# Longitudinal Characterization of a Novel Class of Recurrent, Stereotyped Neurophysiological Events in an Advanced Meditator

**DOI:** 10.64898/2026.07.31.741981

**Authors:** A. L. Callara, F. Bossi, J. Soepa, J. Khechok, N. Sherab, N. Vanello, E. P. Scilingo, B. Neri

## Abstract

Seven EEG recordings over thirteen months revealed highly stereotyped transitions between distinct brain dynamics in an advanced meditator, providing a rare experimental window onto large-scale brain-state dynamics. We report an intensive longitudinal EEG case study of an experienced Tibetan tantric practitioner recorded across seven different measurement sessions, including concentrative and analytical meditation, three Dissolution of Elements sessions, nap, and reading sessions. Across conditions, the EEG repeatedly entered abrupt and reversible “ON” periods lasting tens of seconds. These periods were characterized by high-amplitude delta–theta activity, a structured 7–8 Hz component, fronto-central predominance, and a recurrent transient complex preceding state onset. Compared with matched pre- and post-event intervals, ON periods showed increased spectral power and directed connectivity, reduced relative variability, and high cross-session similarity, consistent with a recurrent and stereotyped macroscopic neurophysiological regime. Additional analyses did not support a straightforward explanation in terms of respiratory-rate changes, sleep-related variations, or overt movement artifacts. During Dissolution of Elements meditation event counts were consistent with the reported structure of the practice. However, given the current state of knowledge about the phenomenon, there is insufficient evidence to establish a close association with either the type of meditation session or the specific practices undertaken during the practitioner’s many years of retreat. We therefore distinguish the robust observation of a recurrent EEG regime from the more tentative hypothesis that it is related to advanced tantric meditative practice. What does appear to be well documented, however, is an unusual, abrupt, and reversible large-scale EEG reconfiguration in a deeply phenotyped expert volunteer, highlighting the value of intensive longitudinal single-participant designs for identifying and characterizing rare neurophysiological phenomena.

## Introduction

Meditation research has undergone a substantial conceptual expansion over the past decades. Initially investigated largely in relation to stress reduction^1^, psychological well-being^2^, and clinical applications^3^, meditation is increasingly recognized as a privileged model for studying attention, emotion regulation, self-related processing, neuroplasticity, and consciousness^4^. Recent theoretical work has emphasized the advent of a “third wave” of meditation research focused on advanced meditation, moving beyond broad categorizations of practice and toward the systematic investigation of meditative states, developmental trajectories, and transformative endpoints that may emerge with sustained training^5^. In this framework, meditation is not only considered as an intervention, but also as an experimental window onto how repeated mental training can shape the dynamics of brain, body, and subjective experience.

A central challenge for this emerging field is that many of the phenomena of interest may be rare, subtle, or difficult to elicit reliably, particularly in practitioners without extensive training. In this context, expert monastic practitioners represent a crucial population for advancing the neuroscience of meditation, because they may be able to reliably enter, sustain, and differentiate refined meditative states under controlled experimental conditions^6^. Prior studies on long-term meditators have shown that extensive contemplative training can be associated with distinctive neural and physiological signatures, including altered oscillatory activity^7^, modulation of attentional and self-referential networks^8^, and practice-specific patterns of brain activation^9^. Moreover, monastic practitioners often train within structured contemplative systems that provide both a stable practice environment and a refined phenomenological vocabulary, facilitating the integration of first-person reports with neurophysiological measures^10,11^.

Beyond cross-sectional comparisons between meditators and controls, repeated-measures designs are particularly well suited to the study of advanced meditation, as they allow state-dependent dynamics to be characterized within the same practitioner across different conditions. This is especially relevant because meditative phenomena are often intrinsically dynamic: they may unfold over seconds or minutes within a session, vary across repeated sessions, or emerge only under specific internal and contextual conditions. Intensive within-subject and neurophenomenological designs can therefore reveal state transitions, intra-individual variability, and practice-specific neural signatures that may be obscured by group-level averaging ^12,13^. This approach is particularly relevant in expert and monastic practitioners, in whom extensive training may permit sufficiently stable engagement with multiple meditative practices to allow direct comparison across conditions ^9,14^. Repeated measurements within the same practitioner can thus open a window onto aspects of meditation that are not readily observable in non-expert samples or in conventional between-group designs, including transient shifts in brain-body dynamics, reproducible markers of meditative depth, and the temporal coupling between subjective experience and neurophysiological reconfiguration.

Empirical investigation of advanced meditative practices remains intrinsically challenging, as practitioners capable of reliably entering highly specialized meditative states are rare and often accessible only through small samples or single-case designs. These limitations are particularly relevant when investigating tantric and other advanced contemplative practices. Because such practices are pursued by only a small number of highly trained individuals and may elicit rare, practice-specific phenomena, standard group-level methodologies can be difficult to apply meaningfully. In these situations, rigorous single-case investigations constitute a valuable early stage of scientific inquiry, thus allowing unusual phenomena to be identified, described, and methodologically characterized. This study addresses this challenge through an intensive case study of a highly experienced monastic practitioner examined across multiple sessions and conditions, within a long-term research program initiated in 2018 through a collaboration between the University of Pisa and the Tibetan Monastic University of Sera Jey^11,15,16^. This program involved the observation of more than 70 meditators, including practitioners with extended retreat experience at Sera Jey Monastery and Gyumed Tantric College, and was supported by prolonged fieldwork inside the monastery in close collaboration with volunteers and the local Science Centre. This sustained engagement was essential for establishing mutual trust and for integrating first-person phenomenological reports with objective neurophysiological measurements, thereby contributing to the still limited neurophenomenological study of advanced meditation in monastic practitioners.

In this study, we report a recurrent and previously uncharacterized EEG phenomenon observed in a highly experienced Tibetan tantric meditator across three ecological measurement campaigns conducted at the participant’s place of residence over a 13-month period. Recordings were obtained under several conditions, including eyes-open and eyes-closed resting state, analytical and concentrative meditation, advanced tantric practices, namely “Dissolution of Elements” meditation, an afternoon nap, and a reading task. Across these conditions, we repeatedly observed the emergence of abrupt and sustained EEG intervals characterized by high-amplitude delta–theta activity and a structured 7-8 Hz component. These episodes typically lasted several tens of seconds, approximately 40–60 s (although longer events were also observed) were consistently preceded by a recurrent transient complex occurring a few seconds before the transition, and showed a broad but non-uniform scalp distribution, with maximal amplitude over frontal regions. Once established, the EEG pattern remained stable for the duration of the episode and then terminated abruptly, with a return to baseline activity. Importantly, these events were not confined to formal meditation, but were also observed during resting state and active cognitive engagement. In the waking conditions, no overt loss of posture or behavioural disruption was observed, although behavioural responsiveness was not assessed continuously on an event-by-event basis. The occurrence across multiple conditions indicates that the phenomenon is not exclusively tied to a specific meditative content or state transition, although its relationship to long-term contemplative training remains to be established.

Nevertheless, while these episodes may occur under various conditions and across different mental states, a highly distinctive temporal organization was observed during sessions of a specific tantric meditation practice known as the *Dissolution of the Elements*. This practice unfolds through a precisely defined sequence of stages, and remarkably, the number of detected events matched the number of transitions between successive stages. This observation raises the possibility that the phenomenon, besides occurring spontaneously, may also be temporally organized or modulated by specific meditative operations. However, in accordance with his monastic commitments, the practitioner did not provide real-time reports of specific contemplative stages. Consequently, individual EEG events could not be directly aligned with the internal progression of the practice.

The primary aim of this study was therefore to characterize the temporal structure, spectral features, spatial distribution, reversibility, and cross-session reproducibility of this recurrent EEG pattern. A secondary, exploratory aim was to assess whether its occurrence showed a distinctive organization during Dissolution of Elements meditation. By documenting this phenomenon in a highly trained practitioner across repeated sessions and experimental conditions, we provide an intensive longitudinal characterization of an unusual and reversible large-scale EEG regime, while distinguishing the robust empirical observation from more tentative interpretations concerning its contemplative origin and functional significance.

## Results

### Longitudinal observation of an expert practitioner

The volunteer, aged 53 years, participated in seven experimental sessions conducted over a period of 13 months. The participant had accumulated more than 15000 hours of meditative practice and had completed several extended retreats, ranging from six months to two years, with a particular focus on advanced tantric practices associated with the completion stage of Tantra, known as Dzogrim.

The experimental protocol included one concentrative meditation session, one analytical meditation session, three sessions involving the Dissolution of Elements meditation, one afternoon nap session, and one reading session (Table S1 in the Supplementary Material). The meditative sessions comprised practices with different attentional and phenomenological demands: Concentrative meditation, based on sustained single-pointed attention; Analytical meditation, involving the detailed analysis of a specific Buddhist teaching concept; and Dissolution of Elements meditation, an advanced Dzogrim practice characterized by a structured progression through increasingly subtle bodily and mental states, traditionally culminating in the Clear Light experience. Detailed descriptions of each practice are provided in Supplementary Section S2.

### Transient state transitions identification: abrupt changes in EEG spectral power, theta-band power envelope, and event distribution

Brain activity during each session was assessed using EEG. Time-resolved power spectral density, spectrograms, and theta-band power envelopes were estimated to characterize temporal changes in neural activity (Figure 1). Across sessions, we observed abrupt transitions into transient states characterized by high-amplitude, low-frequency activity, predominantly within the delta–theta range. These events showed peak-to-peak amplitudes several-fold greater than the ongoing alpha rhythm. In addition to exhibiting a continuous spectral component across the aforementioned frequency bands, this activity was also characterized by a prominent periodic component at approximately 7 Hz. This gave rise to a composite waveform combining large-amplitude slow activity with rhythmic theta-band organization. Importantly, most intervals were preceded by a stereotyped transient complex occurring approximately 1–5 s before event onset (Figure 1a).

**Figure 1.**
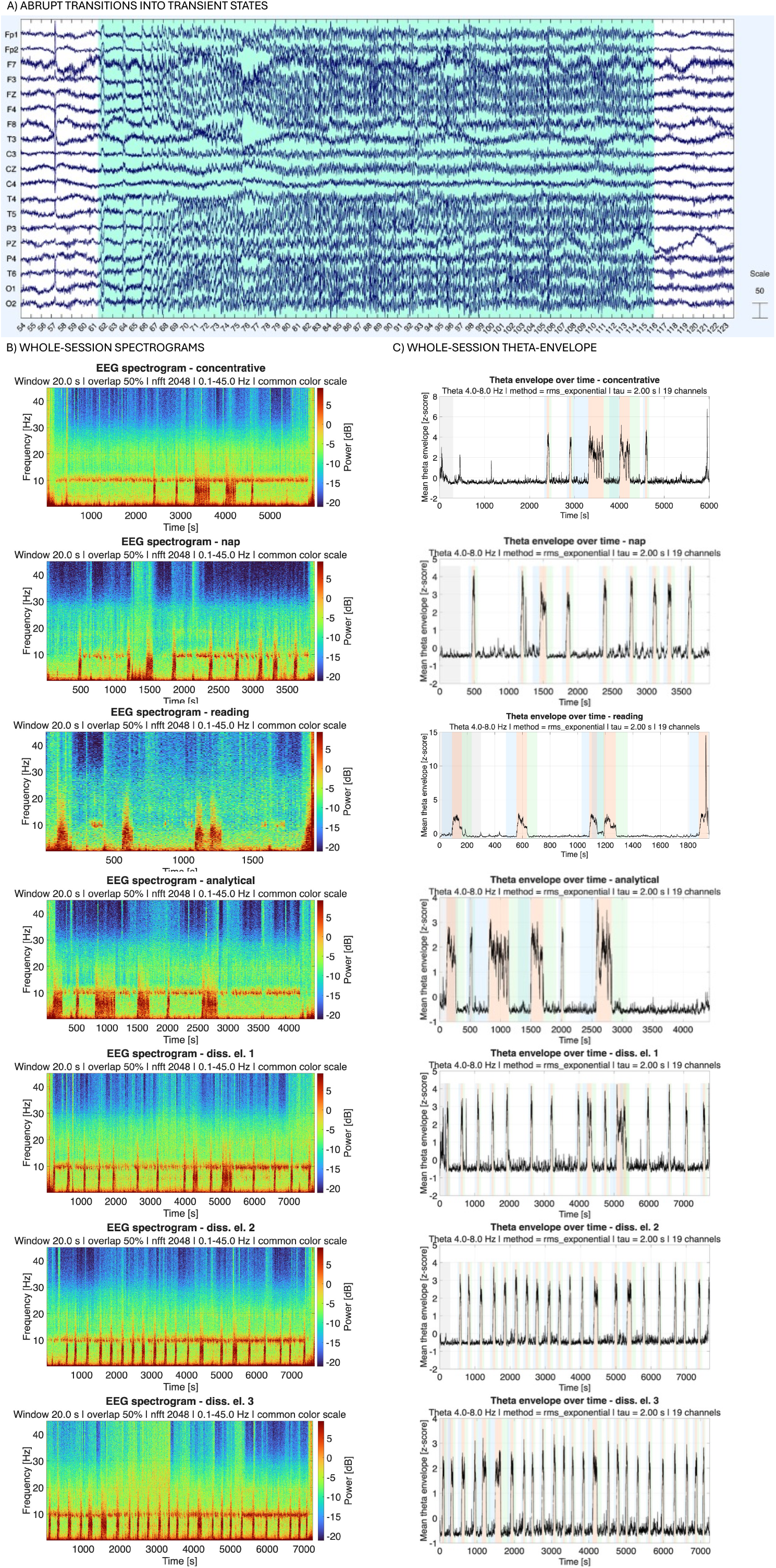
Abrupt transitions into recurrent transient EEG states across experimental sessions. A) Representative EEG segment showing the abrupt onset and offset of a transient high-amplitude state. The ON period is characterized by a sudden increase in low-frequency activity, preceded by a stereotyped transient complex occurring a few seconds before state onset, and followed by a rapid return to the pre-event EEG pattern. B) Whole-session spectrograms for the seven experimental recordings, showing repeated transient increases in low-frequency spectral power across different behavioural and meditative contexts. C) Whole-session theta-band power envelopes, highlighting the temporal stability and recurrence of the detected ON periods across sessions. Shaded intervals indicate detected transient states.

The distribution of events across sessions is summarized in Table 1. Event duration and temporal organization differed across experimental sessions. Analytical and Concentrative meditation were associated with longer-lasting events, whereas Dissolution of Elements meditation showed a more structured and periodic pattern of occurrence. Similar periods, although less structured, were also observed during the afternoon nap and reading sessions.

**Table 1.** Description of transient state duration for each experimental session.

| Session |  | On-period |  |  |  |  |
| --- | --- | --- | --- | --- | --- | --- |
| type | Session duration (s) | Number of events | Mean duration (s) | Median duration (s) | St. dev. (s) | Range (s) |
| Concentrative | 6005 | 5 | 143.40 | 53.00 | 134.55 | 46.00 – 341.00 |
| Nap | 3909 | 9 | 55.89 | 53.00 | 16.47 | 42.00 – 98.00 |
| Reading | 1955 | 5 | 71.80 | 72.00 | 11.32 | 56.00 – 87.00 |
| Analytical | 4438 | 6 | 174.83 | 181.50 | 118.53 | 40.00 – 337.00 |
| Diss. El. #1 | 7745 | 16 | 72.56 | 60.00 | 35.71 | 48.00 – 183.00 |
| Diss. El. #2 | 7665 | 20 | 68.55 | 64.00 | 21.24 | 44.00 – 129.00 |
| Diss. El. #3 | 7241 | 25 | 66.60 | 56.00 | 30.94 | 39.00 – 180.00 |

To assess whether ON-state transitions could be explained by changes in respiratory dynamics, we extracted the respiratory belt signal over the same PRE, ON and POST intervals used for EEG analysis and estimated breathing frequency in each segment. No systematic PRE–ON–POST modulation of breathing frequency was observed across sessions (Supplementary section S6, Supplementary Figure S1). This suggests that the large EEG spectral and connectivity changes observed during ON periods were not driven by concurrent changes in respiratory rate.

Notably, the Dissolution of Elements meditation appeared to be particularly relevant to the organization of these transient state transitions. Specifically, the number of detected transitions appeared to mirror the structured progression of the practice. In our protocol, each Dissolution of Elements session lasted approximately 130 minutes. During each session, the participant repeated the complete sequence twice in both directions, from the grossest element to the subtlest and then in reverse order, as a procedure intended to reinforce the effects of the practice. The full sequence from the Earth element to Clear Light comprised five transitions: Earth to Water, Water to Fire, Fire to Air, Air to Void, and Void to Clear Light. Performed forward and backward and repeated twice, this procedure yielded 20 transitions in total.

Although the practitioner willingly provided general information about the practice, he declined to report, during the recording session, the attainment of any specific meditative state. This limitation reflects his monastic commitments regarding the disclosure of specific contemplative experiences.

### Transient-state transitions: time-dependent changes in EEG spectral power, directed connectivity and scalp topography

Brain activity before, during, and after transient states, hereafter referred to as “ON periods”, was characterized in terms of global spectral power, directed connectivity, and scalp topography. After identifying the ON periods, time-matched segments of equal duration were extracted immediately before and after each event and labelled as PRE and POST, respectively. Changes in EEG power spectral density and directed Direct Transfer Function (dDTF, (Korzeniewska et al., 2008; Mullen, 2010)) were then assessed across the three periods (Figure 2; Supplementary Sections S3 and S4).

**Figure 2.**
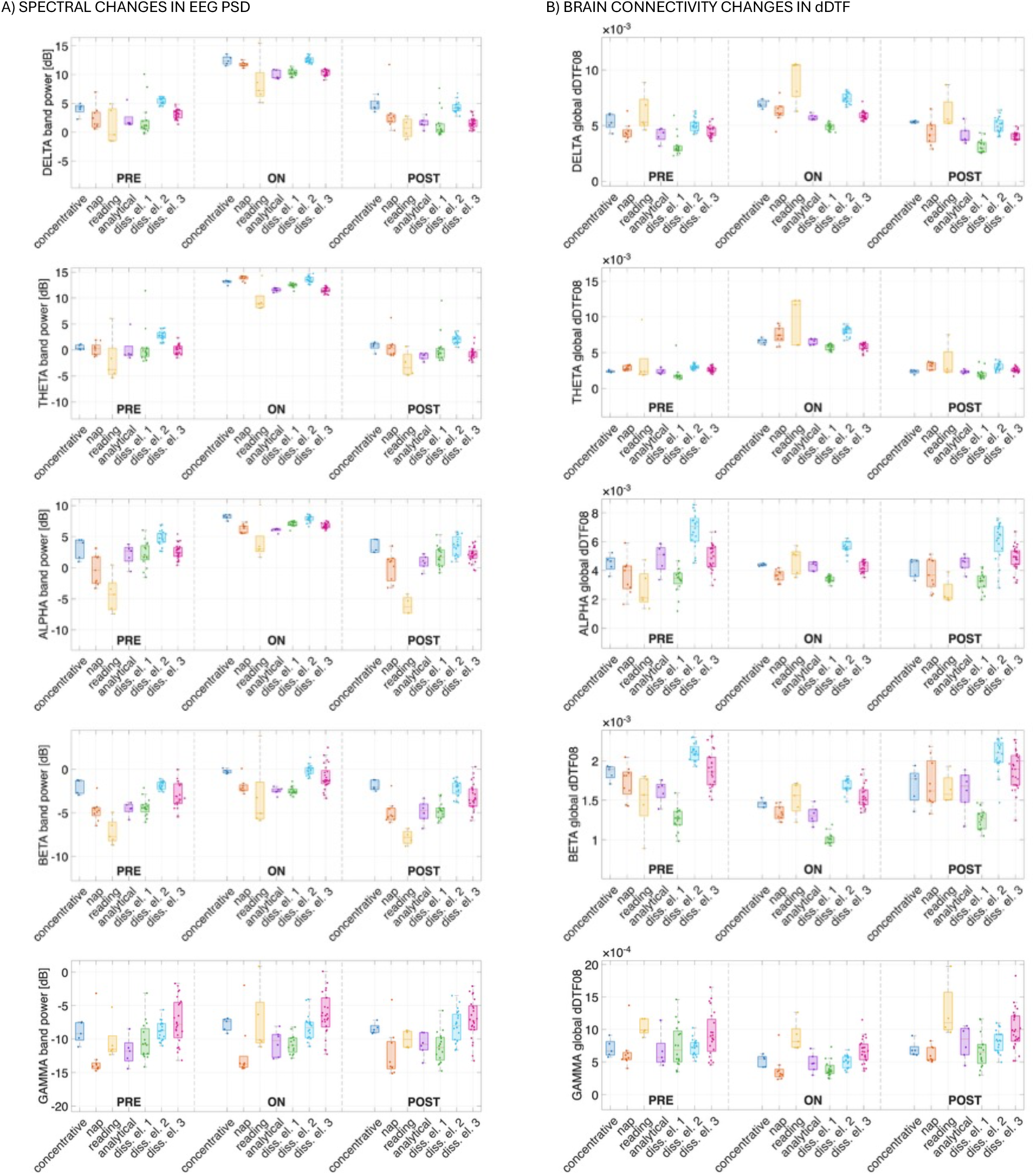
Large-amplitude reversible reconfiguration of EEG power and directed connectivity during ON periods. A) Boxplots of global EEG spectral power in the delta, theta, alpha, beta and gamma bands during matched PRE, ON and POST periods. Across sessions, ON periods were associated with a pronounced elevation of spectral power, particularly in the delta and theta bands, where differences relative to PRE and POST reached the order of ∼10 dB or more. The close overlap between PRE and POST distributions indicates that the transition was reversible and was followed by a restoration of the baseline spectral profile. B) Boxplots of global dDTF connectivity across the same frequency bands and periods. Directed connectivity showed a parallel state-dependent modulation, with ON periods differing from both PRE and POST and with limited PRE–POST differences.

Significant differences in both PSD and dDTF were observed across sessions and frequency bands. Post-hoc analyses showed that PSD changes were most prominent between PRE and ON, and between ON and POST, across nearly all frequency bands, while showing non-significant variations between PRE and POST periods. Gamma-band power represented an exception, showing no significant differences across ON and PRE/POST periods. Global dDTF showed a similar pattern, with differences occurring mainly between PRE and ON, and between ON and POST, across all frequency bands but virtually no change between PRE and POST. Among the experimental conditions, the Dissolution of Elements sessions showed the most pronounced connectivity changes between PRE and ON and between ON and POST.

In contrast to the marked electrophysiological changes, breathing rate showed no systematic variation across matched PRE, ON, and POST periods (Supplementary Material S6; Figure S1).

In addition to the marked increase in mean power and connectivity during ON periods, ON epochs also showed reduced relative dispersion across segments. When variability was expressed as the coefficient of variation, ON periods showed substantially lower relative variability, approximately 30%, compared with PRE and POST periods, where relative variability approached 100%. This pattern suggests that the ON state was not only stronger in magnitude, but also more internally stable and stereotyped across events.

The scalp distribution of spectral changes is shown in the topographical maps in Figure 3. These changes were most prominent over fronto-central regions and were larger at lower frequencies, particularly in the delta and theta bands, decreasing progressively at higher frequencies.

**Figure 3.**
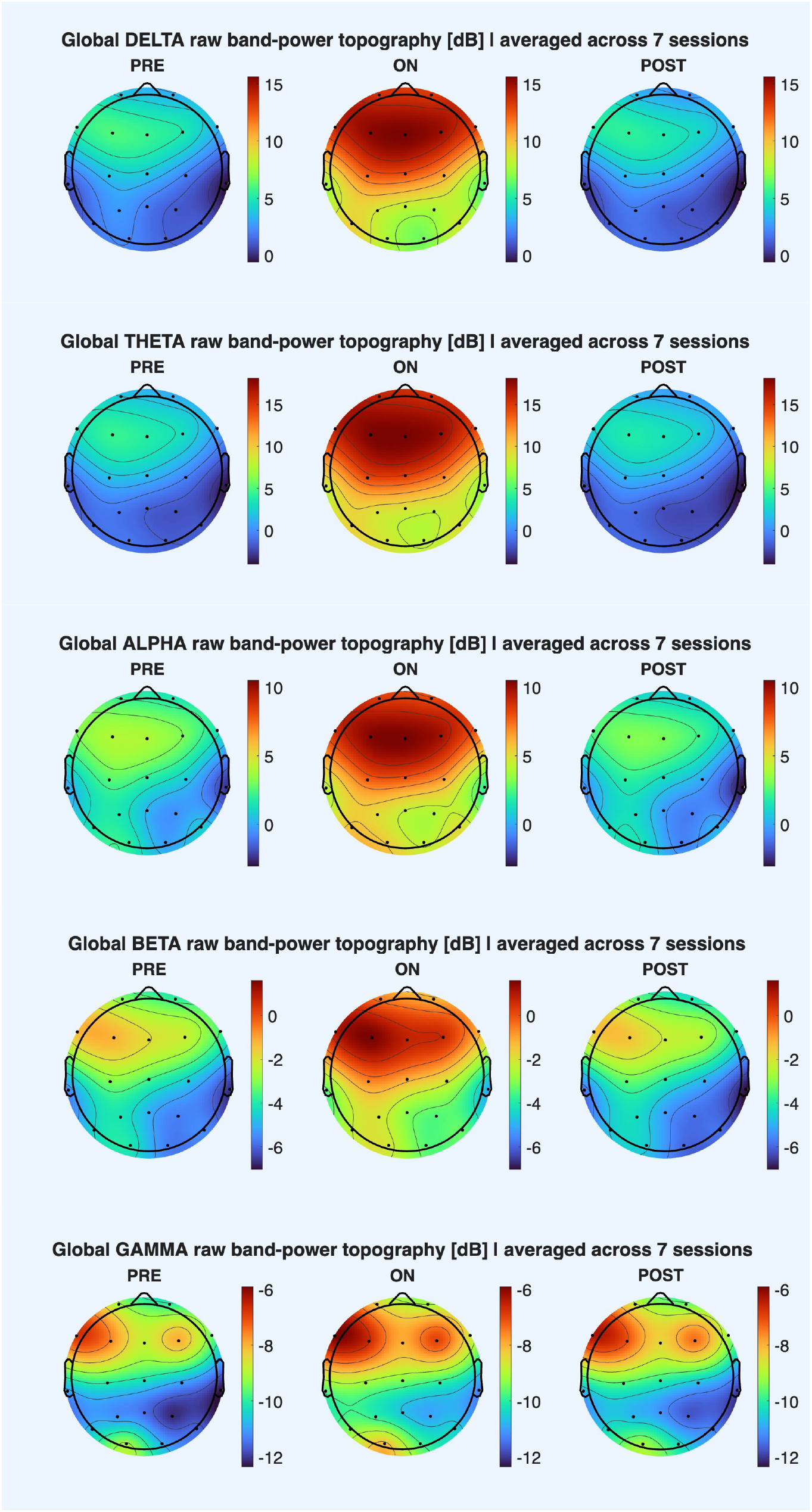
Spatial organization of the transient ON state. Scalp maps display raw band-power topographies, expressed in dB and averaged across the seven sessions, for PRE, ON and POST periods. The ON state produced a large fronto-central power increase, maximal in the delta and theta bands and reaching approximately 10 dB or more relative to PRE and POST. This spatial pattern progressively attenuated in alpha, beta and gamma bands. The similarity between PRE and POST maps indicates that the ON state was spatially and spectrally reversible.

### Cross-session stability of transient-state spectral profiles

The stability of PRE, ON, and POST periods across experimental sessions was assessed using correlation analysis of EEG power spectral density (Figure 4). For each session, average PSD profiles were estimated separately for the PRE, ON, and POST periods. Cross-session correlations were then computed within each condition, comparing PRE with PRE, ON with ON and POST with POST periods across sessions.

**Figure 4.**
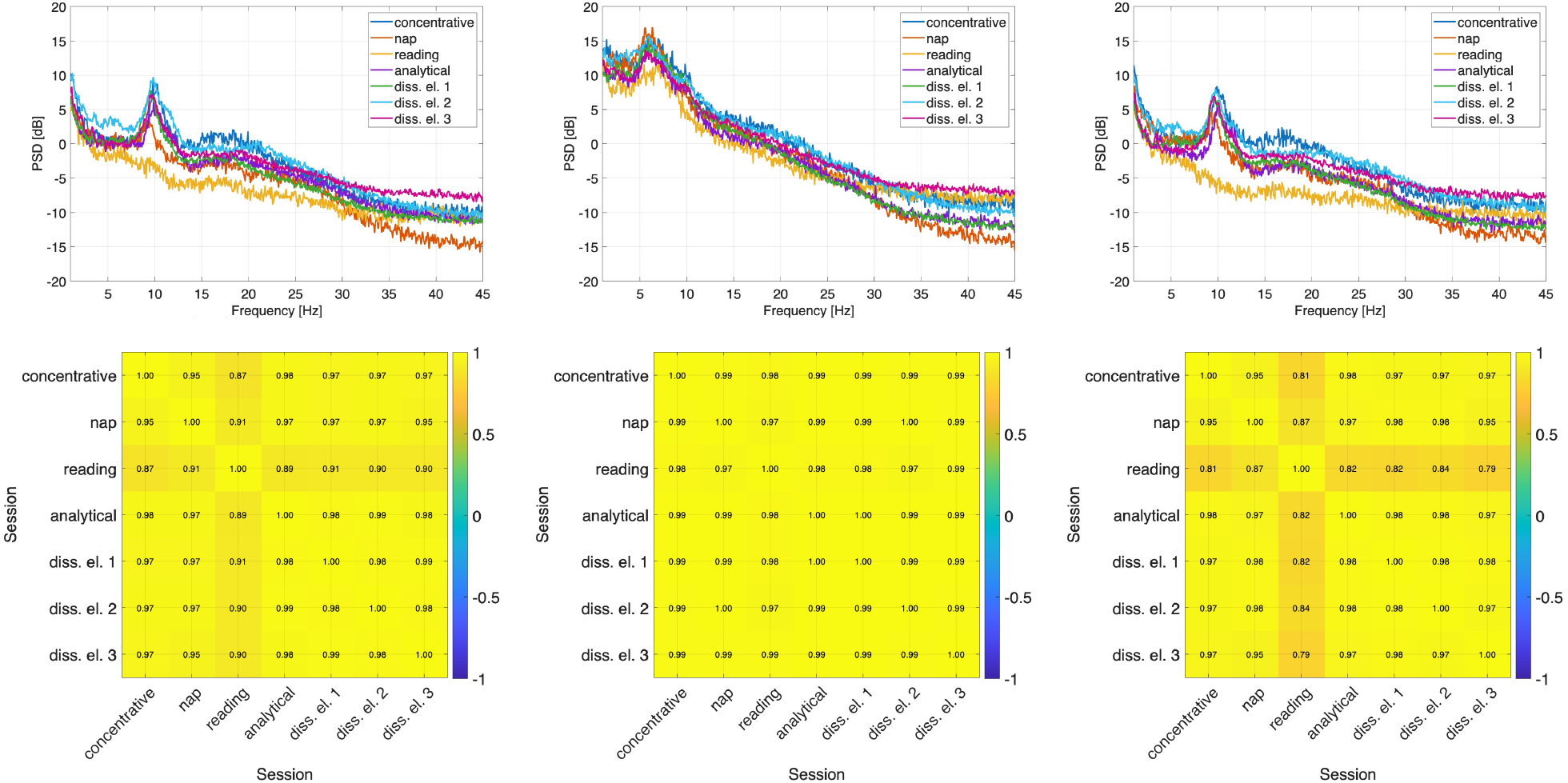
Cross-session stability of PRE, ON and POST spectral profiles. Average Welch PSD profiles were estimated separately for PRE, ON and POST periods in each experimental session and compared across sessions using pairwise correlation matrices. The upper panels show session-level PSD curves for each period, while the lower panels show cross-session correlations between spectral profiles. ON periods displayed highly consistent PSD organization across sessions, with correlation values approaching 1, indicating a reproducible spectral signature of the transient state. PRE and POST periods also showed high cross-session similarity, although with slightly lower consistency than ON periods. The reading session showed comparatively reduced similarity, likely reflecting the absence of a pronounced alpha peak during active cognitive engagement. Overall, the analysis supports the stability of the transient-state spectral profile across the longitudinal protocol.

We observed high cross-session correlations, with values approaching 1, indicating marked stability of the spectral profiles associated with PRE, ON and POST periods across the experimental protocol. The reading session represented the only exception, showing lower correlations with the other sessions, although values remained relatively high, reaching up to 0.81. This lower similarity was mainly attributable to the absence of a clear alpha peak, consistent with the participant’s cognitive engagement during the reading task. Notably, ON periods showed the highest consistency across sessions, exhibiting the strongest correlation values. PRE and POST periods were also highly correlated across sessions, but to a lesser extent than ON periods. Because correlations between full PSD profiles may partly reflect their shared broadband shape, these findings should be interpreted as evidence of cross-session spectral similarity rather than as definitive proof of a unique state-specific signature.

## Discussion

In this intensive longitudinal case report, we describe a novel phenomenon of recurrent, abrupt transitions into a distinct large-scale electrophysiological regime in a long-term meditator. This discrete EEG pattern is defined by high-amplitude delta-theta activity paired with a structured 7–8 Hz theta/low-alpha component, exhibiting a clear fronto-central predominance and a stable duration on the order of tens of seconds. It was reproducibly observed across seven experimental sessions (including resting state, different meditation practices, afternoon nap, and reading) spanning a 13-month period, these transitions display a well-defined ON/OFF temporal architecture that consistently concludes with a complete return to the baseline spectral profile. Our core finding therefore provides evidence that, in this participant, brain activity could repeatedly enter a large-amplitude, reversible, and macroscopically stable EEG configuration without overt behavioural disruption.

This main finding is supported by the identification of a reproducible ON state that differs sharply from both the preceding PRE period and the subsequent POST period across spectral power, functional connectivity, and scalp topography. This convergence across multiple EEG-derived measures supports the interpretation that the ON periods do not reflect a gradual fluctuation around baseline activity, but rather a distinct, highly recurrent, and stereotyped neurophysiological regime. Specifically, post-hoc analyses demonstrated that power spectral density (PSD) and global directed connectivity, estimated via the directed Direct Transfer Function (dDTF), shifted significantly during the ON periods across nearly all frequency bands, while showing non-significant variations between the PRE and POST baselines.

The PRE-ON-POST structure indicates that these events were not readily explained by gradual drifts in vigilance or nonspecific physiological instability, but discrete transitions between relatively stable dynamical regimes. Crucially, correlation analyses of the EEG power spectral density profiles revealed exceptionally high cross-session stability, with correlation values approaching 1. Importantly, the ON state showed higher inter-session similarity than both the PRE and POST periods, indicating that the phenomenon is not simply determined by the global context or specific mental engagement of each individual recording session. Rather, ON epochs appear to share a common, stereotyped spectral organization across the entire 13-month experimental protocol. Because correlations between full PSD profiles can partly reflect their shared broadband spectral shape, this result should be interpreted as evidence of marked cross-session similarity rather than definitive proof of a unique state-specific signature.

To properly contextualize these findings within the current neuroscientific literature, it is essential to distinguish the empirical observation from its subsequent theoretical interpretation. To our knowledge, this study provides a detailed longitudinal characterization of a novel recurrent, macroscopic, and highly structured EEG regime with marked amplitude and prolonged stability. By monitoring the practitioner continuously across multiple resting, meditative, sleep-related, and cognitively demanding conditions over a 13-month longitudinal period, we documented a phenomenon that appears unusual in the combination of its temporal, spectral, and spatial features. The thorough characterization of these parameters constitutes the main empirical contribution of the study and remains distinct from hypotheses concerning the origin or functional significance of the phenomenon.

In contrast to the robust observation of the ON state, interpretations regarding its exact etiology, nature, and systemic implications represent hypotheses rather than definitive conclusions. Mechanistically, the striking coexistence of high-amplitude slow delta-theta activity with a highly structured 7–8 Hz component may suggest a multi-scale coupling between slow excitability fluctuations and thalamo-cortical or fronto-central oscillatory organization. We hypothesize that this distinct configuration may be related to the participant’s multi-decadal contemplative expertise, possibly representing an acquired neurodynamic capacity that may be recruited or modulated by specific advanced tantric practices. However, the present single-case design cannot establish either a causal relationship with contemplative training or the degree of voluntary control over the phenomenon.

From a dynamical systems perspective, these empirical observations may reflect a form of macroscopic metastable switching between large-scale neural configurations. Unlike conventional EEG microstates, which typically unfold and fluctuate at millisecond timescales, the anomalous regimes documented here lasted for tens of seconds (averaging approximately 40-60 s) and involved a stable, large-amplitude reorganization of both spectral power and directed information flow (Fig. 2b). This suggests that, in this participant, wakeful brain activity could access slower and more macroscopically organized EEG regimes than those usually described in conventional resting-state or cognitive EEG paradigms. The presence of a pre-onset transient complex occurring approximately 1-5 s before the transition onset further raises the possibility of a transition-associated precursor. However, its predictive sensitivity and specificity were not formally tested, and it should not yet be interpreted as evidence of a deterministic bifurcation or a transition between distinct attractor-like neural configurations.

Taken together, these findings suggest a dual pattern of the observed phenomenon: cross-condition occurrence and practice-specific organization. On the one hand, the ON state occurred across all recorded baseline and task conditions, including eyes-open/closed rest, an afternoon nap, active reading, and general concentrative or analytical meditation. This cross-condition recurrence is consistent with a relatively stable individual neurophysiological characteristic or predisposition, which may have been influenced by years of intensive contemplative training^5,7,9^. This longitudinal observation was successfully executed within a long-term research program initiated in 2018 to investigate advanced meditation^11,15,16^.

On the other hand, the phenomenon displayed a distinctive temporal organization during sessions dedicated to the Dissolution of the Elements, an advanced completion-stage (Dzogrim) tantric practice. While the ON state was observed across multiple mental and behavioural conditions, its occurrence appeared more structured during this specific practice. Across the three sessions, the number of detected ON episodes was of the same order as the number of transitions expected from the reported structure of the practice. However, because the practitioner did not provide real-time reports of specific contemplative stages, in accordance with his monastic commitments, individual EEG events could not be directly aligned with the internal progression of the practice. These observations raise the possibility that the temporal organization of the ON state may be influenced by specific contemplative operations, although this interpretation remains exploratory.

Given the unusual nature of this high-amplitude delta-theta profile, several alternative physiological and pathological hypotheses were considered. (i) Although the emergence of prominent slow-wave activity raises an obvious comparison with local sleep phenomena during wakefulness^17^ or slow activity reported in other contemplative practices such as Yoga Nidra^18^, several features do not closely support a conventional sleep-related interpretation. In the waking conditions, the episodes were not accompanied by overt loss of posture or observable behavioural responsiveness, and cognitive performance was not assessed continuously. Furthermore, the broad topographical scalp distribution of the delta-theta power (predominantly fronto-central) is inconsistent with the typically localized nature of local sleep events. Finally, the persistence of an organized and structured alpha rhythm during these anomalous intervals is electrophysiologically incompatible with standard NREM sleep dynamics, which require clear alpha suppression. (ii) Abrupt regime transitions are also observed in pathological conditions, most notably in absence epilepsy, where highly structured spike-wave discharges reflect switching between abnormal network states^19^. However, the observed waveforms lacked the characteristic spike-wave morphology typical of epileptiform discharges. Moreover, the events displayed a much broader and more complex frequency structure, were perfectly tolerated behaviourally, and were followed by an organized return to baseline. No overt impairment of awareness was observed, although awareness was not formally assessed during each event. These features make a typical absence-seizure pattern unlikely, but they do not constitute a formal clinical exclusion of epileptiform activity. (iii) Changes in respiratory rate are unlikely to provide a simple explanation for the observed EEG transitions, as no systematic differences in breathing rate were observed across matched PRE, ON, and POST periods (Supplementary Material S6; Figure S1). (iv) A purely artefactual explanation

(e.g., movement, muscle/EMG activity, electrode drifting, or postural adjustments) is highly unlikely. Preprocessing pipelines using Independent Component Analysis (ICA) and semi-automatic classification protocols^20–23^ successfully isolated and eliminated non-brain artefactual sources prior to spectral estimation. Slow eye movements are also unlikely to account for the phenomenon. The ON periods were not characterized by stereotyped EOG-like deflections restricted to frontal electrodes, nor by the polarity and morphology expected for slow ocular movements. In addition, the effect extended beyond the most ocular-sensitive channels and was accompanied by coherent spectral and directed-connectivity changes. Jaw movements or bruxism are unlikely to explain the ON state because such activity would be expected to produce broader-band, high-frequency myogenic contamination, often maximal over temporal or peripheral electrodes, rather than a reproducible low-frequency delta–theta regime with fronto-central predominance. When available, video recordings were visually inspected to assess whether ON periods coincided with overt postural adjustments, repetitive movements, or gross motor activity. No systematic movement pattern temporally locked to ON-state onset or offset was observed. An exemplary video segment is provided as Supplementary Video S1. Finally, the strong cross-session reproducibility, the presence of the 1-5 s pre-onset transient complex, the fronto-central topography, and the associated global directed connectivity changes converge to validate a genuine neural origin.

While providing a compelling proof-of-principle, certain methodological limitations must be acknowledged. First, because this study is based on an intensive single-subject design, the generalizability of these findings across wider populations remains constrained. Second, the lack of concurrent, fine-grained, real-time first-person phenomenological reports in the Dissolution of Elements practice prevents a precise second-by-second alignment between specific subjective transitions and neurophysiological events. Third, the use of standard low-density (19-channel) EEG constrains the spatial resolution, limiting our ability to perform source-level localization or cortical reconstruction analyses.

Future investigations should aim to replicate these observations in other highly trained practitioners using intensive within-subject neurophenomenological designs, which are particularly well suited to characterize state-dependent dynamics^5,6,12,13^, including infra-slow fluctuation (ISF) components, which were recently identified as a specific marker of contemplative activity in advanced tantric meditators^24^.

In summary, the strongest conclusion supported by the present data is the existence of a recurrent, abrupt, and reversible macroscopic EEG state that was not observed in our previous recordings and that showed a highly reproducible temporal, spectral, and topographical organization. A more tentative interpretation is that this state may be facilitated by long-term advanced tantric training and may be recruited in a structured way during the Dissolution of Elements practice. This interpretation is supported by the temporal correspondence between detected ON episodes and the reported architecture of the practice, but it cannot be considered definitive in the absence of real-time phenomenological markers and replication in additional practitioners^6,10^.

## Methods

### Ethical approval and subject recruitment

The study was approved by the Committee on Bioethics of the University of Pisa (Review No. 26/2023). The study procedures were conducted in accordance with local legislation and institutional requirements. Written informed consent was obtained from the participant prior to the involvement in the study. All experimental procedures adhered to the principles of Declaration of Helsinki. The volunteer was recruited from among expert practitioners at Sera Jey Monastery through direct verbal invitation.

### Longitudinal study design

The study included seven EEG acquisitions conducted over 13 months. The initial recordings comprised one concentrative and one analytical meditation session. These sessions revealed unusual EEG phenomena that had not been observed in our previous recordings. To evaluate whether these events could be attributable to artefacts and to guide subsequent experimental design, we conducted an in-depth interview with the volunteer after the first two sessions (Supplementary Section S5, transcript of the volunteer interview). The interview indicated that the practitioner had expertise in advanced tantric practices, including the Dissolution of Elements and Clear Light. He was therefore invited to repeat these practices in subsequent sessions, performed on different occasions and under varying experimental conditions. Additional control sessions were acquired to contextualize the observed activity. Specifically, an afternoon nap session was included to assess potential sleep-related contributions, and a reading task was included to examine whether the observed phenomena reflected state-dependent activity or trait-like spectral features.

The longitudinal protocol comprised seven acquisitions: (i) concentrative meditation, (ii) afternoon nap, (iii) reading, (iv) analytical meditation, and (v–vii) three Dissolution of Elements sessions.

### EEG procedure

Recordings were performed in the meditation room of the Meditation Lab at the Sera Jey Science Center. Before the first session, the volunteer completed a demographic and experiential questionnaire assessing age, years of meditation experience, type of meditation usually practised and average daily practice time^11^. The participant was instructed to perform a single meditation practice according to his expertise, with no predefined constraint on session duration, to preserve ecological validity. Each session was preceded by a 5-min eyes-closed resting-state recording. The onset of the meditation period was verbally indicated by the experimenter.

EEG was acquired using a BE PLUS TLM system (EBNeuro S.p.A., Florence, Italy). Recordings were obtained from 19 scalp electrodes positioned according to the international 10–20 system. Electrode impedances were kept below 20 kΩ using conductive gel. Bilateral mastoid electrodes were also recorded. Signals were sampled at 512 Hz.

Preprocessing was performed using a standardized pipeline implemented with custom MATLAB scripts and EEGLAB functions^20^. Data were first low-pass filtered using an anti-aliasing filter with a cutoff frequency of 57.6 Hz and downsampled to 128 Hz. A high-pass filter with a cutoff frequency of 0.1 Hz was then applied. Bad channels were detected and removed using a correlation-based criterion^21^, and subsequently reconstructed using spherical spline interpolation^20^. Independent component analysis was performed using the Infomax algorithm implemented in EEGLAB^20,25^. Independent components were inspected using a semi-automatic procedure combining ICLabel classification with expert visual inspection^22,23^. Components classified as non-brain sources, including ocular and muscular artefacts, were removed. The EEG signal was then reconstructed from the remaining brain-related components. To exclude possible residual ocular artefacts, ON periods were visually inspected for slow eye-movement signatures, including large frontal deflections with polarity reversal and stereotyped EOG-like morphology. We further inspected the spectra and time courses for broad-band high-frequency activity compatible with muscle or jaw-related artifacts.

### EEG analysis

EEG data were analysed to characterize temporal dynamics, spectral content, global power changes, and directed functional connectivity.

#### Time-course and spectrogram inspection

Preprocessed EEG signals were visually inspected to identify temporal patterns and periods of interest. Particular attention was given to activity in the delta, theta, and alpha frequency bands. Time–frequency representations were computed using the short-time Fourier transform (STFT) with 20-s long windows. Spectrograms were visually examined to identify transient bursts, sustained oscillatory activity, and slow changes in spectral content across the recording. Delta, theta, and alpha activity were specifically considered to characterize state-dependent changes in brain dynamics across the session.

#### Global spectral power

Global spectral power was estimated using a sliding-window approach with non-overlapping 5-min long windows. Within each window, the power spectral density (PSD) was estimated separately for each electrode using Welch’s method with 20-s long segments, 50% overlap, and zero-padding to 2048 samples. Electrode-wise PSDs were then averaged across the scalp to obtain a global PSD estimate for each time window. PSD values were expressed in decibels. This procedure was used to characterize time-varying changes in the distribution of spectral power throughout the recording. Band-limited power was extracted for the delta, theta, alpha, beta, and gamma bands and averaged across electrodes to obtain one global power value per segment and frequency band.

For the statistical comparison of predefined periods of interest, EEG data were segmented into three time intervals around each event: a PRE-event period, an ON-period, and a POST-event period. The duration of the PRE- and POST-event windows was matched to the duration of the corresponding ON-period.

#### Connectivity analysis

Directed functional connectivity was estimated from brain-related independent component time courses using the directed Direct Transfer Function (dDTF), implemented in the SIFT EEGLAB plug-in (dDTF08)^26,27^. A multivariate autoregressive model was estimated in a time-varying manner using 10-s windows with a 10-s step size. Model order was selected as the average optimal order across windows according to the Akaike Information Criterion. Model parameters were estimated using the Vieira–Morf algorithm.

Model adequacy was assessed by evaluating residual whiteness, model consistency, and model stability. Residual whiteness was used to assess the absence of residual autocorrelation. Model consistency quantified the proportion of the data correlation structure explained by the model. Model stability was used to verify that the estimated autoregressive process remained stationary.

The estimated autoregressive coefficients were then used to compute dDTF, defined as the product of the full-frequency Direct Transfer Function and partial coherence. dDTF provides a frequency-domain measure of directed, conditionally dependent information flow and can be interpreted in relation to conditional Granger-causal interactions.

For each session and frequency band, a global connectivity value was obtained by averaging dDTF values across all directed connections, excluding self-connections, across the selected frequency bins, and across time windows falling within each PRE-, ON-, and POST-period. This yielded one global connectivity estimate per segment, condition, session, and frequency band.

#### Statistical analysis

Statistical analyses were performed separately for global spectral power and global directed connectivity. For each frequency band and session, PRE-, ON-, and POST-period values were treated as repeated measures across matched segments. Condition effects were assessed using repeated-measures analysis of variance.

Because tests were performed across multiple frequency bands and sessions, p-values from the global condition tests were corrected across all session-by-band comparisons using the Benjamini–Hochberg false discovery rate procedure^28^. Thus, for five frequency bands and seven sessions, correction was applied across 35 global tests.

When a global condition effect was identified, post-hoc paired comparisons were performed between PRE and ON, PRE and POST, and ON and POST periods. Post-hoc comparisons were conducted using paired t-tests. Post-hoc p-values were corrected within each session and frequency band across the three pairwise comparisons using Holm correction. Effect sizes for paired comparisons were quantified using Cohen’s dz, computed as the mean paired difference divided by the standard deviation of the paired differences.

Statistical significance was assessed at an adjusted threshold of α = 0.05.

## Declaration of generative AI and AI-assisted technologies in the writing process

During the preparation of this work the authors used ChatGPT 5.1 and GEMINI Flash 2.5 in order to improve the readability and language of the manuscript. After using these tools/services, the authors reviewed and edited the content as needed and take full responsibility for the content of the published article.

