## Supplementary Material for "Longitudinal Characterization of a Novel Class of Recurrent, Stereotyped Neurophysiological Events in an Advanced Meditator"

**Supplementary materials**

**S1. Description of each experimental session.**

Table 1. Description of each experimental session.

| Session | Description | Date |
| --- | --- | --- |
| 1. Concentrative | 5 minutes of eyes-closed resting state followed by a concentrative session whose duration was up to volunteers choice. | Jan 2025 |
| 2. Nap | 10 minutes of eyes-closed resting state followed by a nap session whose duration was up to volunteers choice. | May 2025 |
| 3. Reading | 5 minutes of eyes-open resting state followed by a reading session. | Feb 2025 |
| 4. Analytical | 5 minutes of eyes-closed resting state followed by an analytical session whose duration was up to volunteers choice. | Jan 2025 |
| 5. Dissolution of Elements 1 | 5 minutes of eyes-closed resting state followed by dissolution of elements session whose duration was up to volunteers choice. | April 2025 |
| 6. Dissolution of Elements 2 | 5 minutes of eyes-closed resting state followed by dissolution of elements session whose duration was up to volunteers choice. | April 2025 |
| 7. Dissolution of Elements 3 | 5 minutes of eyes-closed resting state followed by dissolution of elements session whose duration was up to volunteers choice. | Feb 2026 |

**S2. Description of meditative practices**

*Concentrative meditation:* In this practice, the meditator focuses the mind undistractedly on a single point, such as a visualized image (e.g., the Buddha), the natural flow of the breath, or the recitation of a mantra, without analytical elaboration. This meditation develops the ability to focus and serves as the basis for other types of practice. The intention is to cultivate a state of wakefulness that is neither fluctuating nor dull, working to pacify the two main obstacles to concentration: laxity and excitement, to achieve the “factor of clarity” and “factor of stability.”

*Analytical meditation:* analytical meditation is a form of meditation in which the meditator initially directs his/her attention to what appears to the mind without attachment and by cultivating awareness. Then, the meditator focuses and analyzes in depth a particular concept of the Dharma (e.g., emptiness, compassion, lamrim, consciousness and its levels). As a result, this practice corresponds to a clear stabilized mind and is well described as a logic/conceptual task.

*Dissolution of elements:* The Dissolution of the Elements and Clear Light meditation belongs to the completion stage (Dzogrim) of Highest Yoga Tantra in the Tibetan Buddhist tradition. It is a structured contemplative process that models, in a controlled and reversible manner, the inner sequence traditionally described as occurring at the time of death. The practice consists of intentionally generating progressively subtler states of consciousness, beginning with the ordinary or gross state, characterized by the predominance of bodily processes (the sensory consciousnesses), and proceeding through increasingly subtle levels until reaching a state known as the Clear Light Consciousness, also referred to as the primordial or extremely subtle consciousness. Each state is associated with a different “element”, beginning with the grossest element (earth) and progressing gradually through water, fire, air, and finally space. As the coarse levels of mind subside, subtler layers of consciousness emerge. The Clear Light (Ösel) consciousness is described as a state of luminous, non-dual awareness devoid of conceptual elaboration. In the Tibetan Buddhist tradition, it is said that the Clear Light mind state, in addition to manifesting at the moment of death, is also experienced during life by ordinary individuals (who, however, do not retain awareness of it) during phases of deep, dreamless sleep. Advanced practitioners, by contrast, due to their exceptionally high level of training, are able to experience it with full awareness, both during deep sleep and in specific contemplative practices such as Dzogrim. Therefore, Clear Light is not mere blankness or sleep-like quiescence; it is characterized as reflexive, self-knowing clarity, void of object, free from discursive thought yet vividly present. Advanced practitioners train to recognize and stabilize this state during meditation, with the soteriological aim of realizing emptiness directly and preparing for the death process. In traditional doctrine, mastery of Clear Light practice is said to transform death into an opportunity for liberation.

**Supplementary Table S3. Repeated-measures ANOVA and post-hoc comparisons of EEG spectral power across PRE, ON and POST periods**

============================================================
GLOBAL PSD PRE / ON / POST TESTS ACROSS ALL BANDS AND SESSIONS
============================================================

| **Analysis** | **BandName** | **BandLowHz** | **BandHighHz** | **SessionIndex** | **SessionName** | **NPairedChunks** | **GlobalTest** | **Statistic** | **DF1** | **DF2** | **PValue** | **PAdjusted** | **Significant** |
| --- | --- | --- | --- | --- | --- | --- | --- | --- | --- | --- | --- | --- | --- |
| **PSD** | alpha | 8 | 13 | 1 | concentrative | 5 | rm_anova | 42.5818225023066 | 2 | 8 | 5.43718437933774e-05 | 8.65006605803732e-05 | TRUE |
| **PSD** | alpha | 8 | 13 | 2 | nap | 9 | rm_anova | 40.2786390833528 | 2 | 16 | 5.68433599973648e-07 | 1.24344849994235e-06 | TRUE |
| **PSD** | alpha | 8 | 13 | 3 | reading | 4 | rm_anova | 16.8202125790401 | 2 | 6 | 0.00346767861137313 | 0.00466802889992536 | TRUE |
| **PSD** | alpha | 8 | 13 | 4 | analytical | 5 | rm_anova | 45.6134329068658 | 2 | 8 | 4.22515701000951e-05 | 7.04192835001585e-05 | TRUE |
| **PSD** | alpha | 8 | 13 | 5 | diss. el. 1 | 16 | rm_anova | 79.7640692353034 | 2 | 30 | 9.81100202707122e-13 | 3.12168246315903e-12 | TRUE |
| **PSD** | alpha | 8 | 13 | 6 | diss. el. 2 | 20 | rm_anova | 91.6841432071304 | 2 | 38 | 2.87538282201285e-15 | 1.25797998463062e-14 | TRUE |
| **PSD** | alpha | 8 | 13 | 7 | diss. el. 3 | 24 | rm_anova | 221.410854620352 | 2 | 46 | 2.4715475335311e-24 | 1.73008327347177e-23 | TRUE |
| **PSD** | beta | 13 | 30 | 1 | concentrative | 5 | rm_anova | 22.9883406366217 | 2 | 8 | 0.000482542125922 | 0.000734303235098696 | TRUE |
| **PSD** | beta | 13 | 30 | 2 | nap | 9 | rm_anova | 20.5204097524896 | 2 | 16 | 3.8324262845538e-05 | 6.70674599796915e-05 | TRUE |
| **PSD** | beta | 13 | 30 | 3 | reading | 4 | rm_anova | 18.5074557351509 | 2 | 6 | 0.00271391755909673 | 0.00395779644034939 | TRUE |
| **PSD** | beta | 13 | 30 | 4 | analytical | 5 | rm_anova | 12.8124194513781 | 2 | 8 | 0.00320419618320178 | 0.0044858746564825 | TRUE |
| **PSD** | beta | 13 | 30 | 5 | diss. el. 1 | 16 | rm_anova | 52.9040701824276 | 2 | 30 | 1.45523048687897e-10 | 3.63807621719743e-10 | TRUE |
| **PSD** | beta | 13 | 30 | 6 | diss. el. 2 | 20 | rm_anova | 146.090128415756 | 2 | 38 | 1.4441669986202e-18 | 8.42430749195119e-18 | TRUE |
| **PSD** | beta | 13 | 30 | 7 | diss. el. 3 | 24 | rm_anova | 87.4059038528458 | 2 | 46 | 2.14248300562873e-16 | 1.07124150281436e-15 | TRUE |
| **PSD** | delta | 1 | 4 | 1 | concentrative | 5 | rm_anova | 138.496980925064 | 2 | 8 | 6.20893785493404e-07 | 1.27831073483936e-06 | TRUE |
| **PSD** | delta | 1 | 4 | 2 | nap | 9 | rm_anova | 42.0640634501015 | 2 | 16 | 4.25119597806716e-07 | 9.91945728215671e-07 | TRUE |
| **PSD** | delta | 1 | 4 | 3 | reading | 4 | rm_anova | 10.1588769998117 | 2 | 6 | 0.0118496979914283 | 0.0148121224892854 | TRUE |
| **PSD** | delta | 1 | 4 | 4 | analytical | 5 | rm_anova | 72.7245270366788 | 2 | 8 | 7.38759511726345e-06 | 1.36087278475906e-05 | TRUE |
| **PSD** | delta | 1 | 4 | 5 | diss. el. 1 | 16 | rm_anova | 78.5247712009266 | 2 | 30 | 1.195278395164e-12 | 3.48622865256168e-12 | TRUE |
| **PSD** | delta | 1 | 4 | 6 | diss. el. 2 | 20 | rm_anova | 1113.30136109691 | 2 | 38 | 1.86655099778431e-34 | 1.63323212306127e-33 | TRUE |
| **PSD** | delta | 1 | 4 | 7 | diss. el. 3 | 24 | rm_anova | 929.183460646234 | 2 | 46 | 6.44415023786423e-38 | 7.51817527750827e-37 | TRUE |
| **PSD** | gamma | 30 | 45 | 1 | concentrative | 5 | rm_anova | 1.47687349347886 | 2 | 8 | 0.284517593700067 | 0.331937192650078 | FALSE |
| **PSD** | gamma | 30 | 45 | 2 | nap | 9 | rm_anova | 0.204611816068788 | 2 | 16 | 0.81706308743755 | 0.841094354715125 | FALSE |
| **PSD** | gamma | 30 | 45 | 3 | reading | 4 | rm_anova | 0.165121557800266 | 2 | 6 | 0.851515548228922 | 0.851515548228922 | FALSE |
| **PSD** | gamma | 30 | 45 | 4 | analytical | 5 | rm_anova | 1.19140121747816 | 2 | 8 | 0.352453304547169 | 0.385495801848467 | FALSE |
| **PSD** | gamma | 30 | 45 | 5 | diss. el. 1 | 16 | rm_anova | 0.881602778430825 | 2 | 30 | 0.424574535753794 | 0.450306325799479 | FALSE |
| **PSD** | gamma | 30 | 45 | 6 | diss. el. 2 | 20 | rm_anova | 1.13960075301086 | 2 | 38 | 0.330637522278887 | 0.373300428379389 | FALSE |
| **PSD** | gamma | 30 | 45 | 7 | diss. el. 3 | 24 | rm_anova | 2.78147381598024 | 2 | 46 | 0.0723872699079988 | 0.0873639464406882 | FALSE |
| **PSD** | theta | 4 | 8 | 1 | concentrative | 5 | rm_anova | 1189.4483546522 | 2 | 8 | 1.26190146644991e-10 | 3.39742702505744e-10 | TRUE |
| **PSD** | theta | 4 | 8 | 2 | nap | 9 | rm_anova | 256.791811921904 | 2 | 16 | 6.94198202166051e-13 | 2.42969370758118e-12 | TRUE |
| **PSD** | theta | 4 | 8 | 3 | reading | 4 | rm_anova | 11.6920604380991 | 2 | 6 | 0.00851364712509849 | 0.0110362092362388 | TRUE |
| **PSD** | theta | 4 | 8 | 4 | analytical | 5 | rm_anova | 102.398206641327 | 2 | 8 | 1.99757334200093e-06 | 3.88417038722403e-06 | TRUE |
| **PSD** | theta | 4 | 8 | 5 | diss. el. 1 | 16 | rm_anova | 120.598045101799 | 2 | 30 | 4.54535135380505e-15 | 1.76763663759085e-14 | TRUE |
| **PSD** | theta | 4 | 8 | 6 | diss. el. 2 | 20 | rm_anova | 2335.26911807518 | 2 | 38 | 1.70236392370095e-40 | 2.97913686647667e-39 | TRUE |
| **PSD** | theta | 4 | 8 | 7 | diss. el. 3 | 24 | rm_anova | 1871.95421304861 | 2 | 46 | 8.60933586767604e-45 | 3.01326755368661e-43 | TRUE |

============================================================
PSD POST-HOC TESTS ACROSS ALL BANDS AND SESSIONS
============================================================

| **Analysis** | **BandName** | **BandLowHz** | **BandHighHz** | **SessionIndex** | **SessionName** | **NPairedChunks** | **Condition1** | **Condition2** | **Mean1** | **Mean2** | **Test** | **TestStatistic** | **DF** | **CohensDz** | **PValue** | **PAdjusted** | **Significant** |
| --- | --- | --- | --- | --- | --- | --- | --- | --- | --- | --- | --- | --- | --- | --- | --- | --- | --- |
| **PSD** | alpha | 8 | 13 | 1 | concentrative | 5 | PRE | ON | 3.11746613021703 | 8.20361263046232 | paired_ttest | -6.17567868178779 | 4 | 2.76184746793476 | 0.00349196843886134 | 0.00698393687772269 | TRUE |
| **PSD** | alpha | 8 | 13 | 1 | concentrative | 5 | PRE | POST | 3.11746613021703 | 3.37314845934908 | paired_ttest | -0.493575621262306 | 4 | 0.220733728235842 | 0.647478431835512 | 0.647478431835512 | FALSE |
| **PSD** | alpha | 8 | 13 | 1 | concentrative | 5 | ON | POST | 8.20361263046232 | 3.37314845934908 | paired_ttest | 10.5312407984153 | 4 | -4.70971406253517 | 0.000459799235725735 | 0.0013793977071772 | TRUE |
| **PSD** | alpha | 8 | 13 | 2 | nap | 9 | PRE | ON | -0.300606587702012 | 6.25366392840092 | paired_ttest | -8.06986867672416 | 8 | 2.68995622557472 | 4.10164375262882e-05 | 8.20328750525764e-05 | TRUE |
| **PSD** | alpha | 8 | 13 | 2 | nap | 9 | PRE | POST | -0.300606587702012 | 0.0772284603985066 | paired_ttest | -0.379635451603391 | 8 | 0.126545150534464 | 0.714095471327128 | 0.714095471327128 | FALSE |
| **PSD** | alpha | 8 | 13 | 2 | nap | 9 | ON | POST | 6.25366392840092 | 0.0772284603985066 | paired_ttest | 10.1954591542661 | 8 | -3.39848638475537 | 7.34463299609074e-06 | 2.20338989882722e-05 | TRUE |
| **PSD** | alpha | 8 | 13 | 3 | reading | 4 | PRE | ON | -3.68376555868069 | 2.74377874229668 | paired_ttest | -3.3957166782537 | 3 | 1.69785833912685 | 0.0425952662643703 | 0.0851905325287406 | FALSE |
| **PSD** | alpha | 8 | 13 | 3 | reading | 4 | PRE | POST | -3.68376555868069 | -6.03682756924603 | paired_ttest | 1.27382222238935 | 3 | -0.636911111194675 | 0.292456367742087 | 0.292456367742087 | FALSE |
| **PSD** | alpha | 8 | 13 | 3 | reading | 4 | ON | POST | 2.74377874229668 | -6.03682756924603 | paired_ttest | 14.3594896639902 | 3 | -7.17974483199509 | 0.000732019779110965 | 0.00219605933733289 | TRUE |
| **PSD** | alpha | 8 | 13 | 4 | analytical | 5 | PRE | ON | 2.09748342736143 | 6.02390763522427 | paired_ttest | -6.03396686011642 | 4 | 2.69847201464026 | 0.00380299352167976 | 0.00760598704335952 | TRUE |
| **PSD** | alpha | 8 | 13 | 4 | analytical | 5 | PRE | POST | 2.09748342736143 | 0.952004196616246 | paired_ttest | 2.06004503606103 | 4 | -0.921280147468696 | 0.10844700274269 | 0.10844700274269 | FALSE |
| **PSD** | alpha | 8 | 13 | 4 | analytical | 5 | ON | POST | 6.02390763522427 | 0.952004196616246 | paired_ttest | 11.395609006517 | 4 | -5.09627127671616 | 0.000338242473124556 | 0.00101472741937367 | TRUE |
| **PSD** | alpha | 8 | 13 | 5 | diss. el. 1 | 16 | PRE | ON | 2.19151757967391 | 7.08331960219857 | paired_ttest | -9.4173581448168 | 15 | 2.3543395362042 | 1.09427637594306e-07 | 2.18855275188613e-07 | TRUE |
| **PSD** | alpha | 8 | 13 | 5 | diss. el. 1 | 16 | PRE | POST | 2.19151757967391 | 1.79284918155405 | paired_ttest | 0.902287833082508 | 15 | -0.225571958270627 | 0.381164652667432 | 0.381164652667432 | FALSE |
| **PSD** | alpha | 8 | 13 | 5 | diss. el. 1 | 16 | ON | POST | 7.08331960219857 | 1.79284918155405 | paired_ttest | 12.2064381739446 | 15 | -3.05160954348616 | 3.4226609176384e-09 | 1.02679827529152e-08 | TRUE |
| **PSD** | alpha | 8 | 13 | 6 | diss. el. 2 | 20 | PRE | ON | 4.8044128013523 | 7.86750870657986 | paired_ttest | -12.5133722208131 | 19 | 2.79807509134956 | 1.27286819326952e-10 | 2.54573638653904e-10 | TRUE |
| **PSD** | alpha | 8 | 13 | 6 | diss. el. 2 | 20 | PRE | POST | 4.8044128013523 | 3.47608379043519 | paired_ttest | 3.32684692578849 | 19 | -0.743905587679927 | 0.00354425459973882 | 0.00354425459973882 | TRUE |
| **PSD** | alpha | 8 | 13 | 6 | diss. el. 2 | 20 | ON | POST | 7.86750870657986 | 3.47608379043519 | paired_ttest | 13.0882234238758 | 19 | -2.92661572804912 | 5.90039314709876e-11 | 1.77011794412963e-10 | TRUE |
| **PSD** | alpha | 8 | 13 | 7 | diss. el. 3 | 24 | PRE | ON | 2.58162315155525 | 6.74177873190161 | paired_ttest | -19.1755119573897 | 23 | 3.91418498768685 | 1.20814483320075e-15 | 2.4162896664015e-15 | TRUE |
| **PSD** | alpha | 8 | 13 | 7 | diss. el. 3 | 24 | PRE | POST | 2.58162315155525 | 2.08042429527497 | paired_ttest | 1.7860297098886 | 23 | -0.364571787889845 | 0.0872866355432029 | 0.0872866355432029 | FALSE |
| **PSD** | alpha | 8 | 13 | 7 | diss. el. 3 | 24 | ON | POST | 6.74177873190161 | 2.08042429527497 | paired_ttest | 20.5165416497036 | 23 | -4.18792152736106 | 2.77163069752933e-16 | 8.31489209258799e-16 | TRUE |
| **PSD** | beta | 13 | 30 | 1 | concentrative | 5 | PRE | ON | -1.91178709218494 | -0.240066414262912 | paired_ttest | -4.49407582761474 | 4 | 2.00981180931704 | 0.0108717832079141 | 0.0217435664158282 | TRUE |
| **PSD** | beta | 13 | 30 | 1 | concentrative | 5 | PRE | POST | -1.91178709218494 | -1.86810766347604 | paired_ttest | -0.192200127441836 | 4 | 0.0859545100488134 | 0.856948633974373 | 0.856948633974373 | FALSE |
| **PSD** | beta | 13 | 30 | 1 | concentrative | 5 | ON | POST | -0.240066414262912 | -1.86810766347604 | paired_ttest | 7.51512488770126 | 4 | -3.3608660216601 | 0.0016780491306723 | 0.0050341473920169 | TRUE |
| **PSD** | beta | 13 | 30 | 2 | nap | 9 | PRE | ON | -4.74111321783918 | -1.99686271955425 | paired_ttest | -10.3555862813146 | 8 | 3.45186209377154 | 6.53526856703901e-06 | 1.9605805701117e-05 | TRUE |
| **PSD** | beta | 13 | 30 | 2 | nap | 9 | PRE | POST | -4.74111321783918 | -4.94824887229134 | paired_ttest | 0.298833136032867 | 8 | -0.099611045344289 | 0.772682792036909 | 0.772682792036909 | FALSE |
| **PSD** | beta | 13 | 30 | 2 | nap | 9 | ON | POST | -1.99686271955425 | -4.94824887229134 | paired_ttest | 5.9884806406345 | 8 | -1.99616021354483 | 0.000327574163094185 | 0.000655148326188369 | TRUE |
| **PSD** | beta | 13 | 30 | 3 | reading | 4 | PRE | ON | -7.03757757133507 | -4.9602239557353 | paired_ttest | -3.48735167914937 | 3 | 1.74367583957468 | 0.0398429562137782 | 0.0796859124275563 | FALSE |
| **PSD** | beta | 13 | 30 | 3 | reading | 4 | PRE | POST | -7.03757757133507 | -7.82510986186931 | paired_ttest | 1.86845803045588 | 3 | -0.934229015227938 | 0.158495398848697 | 0.158495398848697 | FALSE |
| **PSD** | beta | 13 | 30 | 3 | reading | 4 | ON | POST | -4.9602239557353 | -7.82510986186931 | paired_ttest | 6.7968028664788 | 3 | -3.3984014332394 | 0.00651189754304799 | 0.019535692629144 | TRUE |
| **PSD** | beta | 13 | 30 | 4 | analytical | 5 | PRE | ON | -4.56314013795434 | -2.5476498436061 | paired_ttest | -5.41539570803293 | 4 | 2.42183858564445 | 0.0056342161026452 | 0.0169026483079356 | TRUE |
| **PSD** | beta | 13 | 30 | 4 | analytical | 5 | PRE | POST | -4.56314013795434 | -4.85806407252133 | paired_ttest | 0.840500057643278 | 4 | -0.375883052796572 | 0.447934032224635 | 0.447934032224635 | FALSE |
| **PSD** | beta | 13 | 30 | 4 | analytical | 5 | ON | POST | -2.5476498436061 | -4.85806407252133 | paired_ttest | 3.33887507988716 | 4 | -1.49319032940154 | 0.0288659841439688 | 0.0577319682879375 | FALSE |
| **PSD** | beta | 13 | 30 | 5 | diss. el. 1 | 16 | PRE | ON | -4.38981704131514 | -2.48013654081161 | paired_ttest | -8.02692647890033 | 15 | 2.00673161972508 | 8.26165547747995e-07 | 1.65233109549599e-06 | TRUE |
| **PSD** | beta | 13 | 30 | 5 | diss. el. 1 | 16 | PRE | POST | -4.38981704131514 | -4.79833469210974 | paired_ttest | 1.45304348943288 | 15 | -0.363260872358221 | 0.166811001950997 | 0.166811001950997 | FALSE |
| **PSD** | beta | 13 | 30 | 5 | diss. el. 1 | 16 | ON | POST | -2.48013654081161 | -4.79833469210974 | paired_ttest | 11.8881635675304 | 15 | -2.97204089188259 | 4.91235849668159e-09 | 1.47370754900448e-08 | TRUE |
| **PSD** | beta | 13 | 30 | 6 | diss. el. 2 | 20 | PRE | ON | -1.880712696015 | -0.0875139501791119 | paired_ttest | -21.3457384030067 | 19 | 4.77305220990508 | 9.71756409463516e-15 | 2.91526922839055e-14 | TRUE |
| **PSD** | beta | 13 | 30 | 6 | diss. el. 2 | 20 | PRE | POST | -1.880712696015 | -2.20198479309728 | paired_ttest | 2.09864984458358 | 19 | -0.469272371345825 | 0.0494464243940598 | 0.0494464243940598 | TRUE |
| **PSD** | beta | 13 | 30 | 6 | diss. el. 2 | 20 | ON | POST | -0.0875139501791119 | -2.20198479309728 | paired_ttest | 13.9944231575346 | 19 | -3.12924814861447 | 1.8533730279984e-11 | 3.70674605599681e-11 | TRUE |
| **PSD** | beta | 13 | 30 | 7 | diss. el. 3 | 24 | PRE | ON | -2.97901900542043 | -0.891669337434083 | paired_ttest | -16.0358178065044 | 23 | 3.27329760284769 | 5.60197638500746e-14 | 1.68059291550224e-13 | TRUE |
| **PSD** | beta | 13 | 30 | 7 | diss. el. 3 | 24 | PRE | POST | -2.97901900542043 | -3.18143994595551 | paired_ttest | 0.951487239222954 | 23 | -0.194221519405476 | 0.351249741353642 | 0.351249741353642 | FALSE |
| **PSD** | beta | 13 | 30 | 7 | diss. el. 3 | 24 | ON | POST | -0.891669337434083 | -3.18143994595551 | paired_ttest | 10.4405230099369 | 23 | -2.13116283517768 | 3.36845447698029e-10 | 6.73690895396059e-10 | TRUE |
| **PSD** | delta | 1 | 4 | 1 | concentrative | 5 | PRE | ON | 3.97557802270321 | 12.4489794168014 | paired_ttest | -12.7409548123779 | 4 | 5.69792821174599 | 0.000218632613286849 | 0.000437265226573698 | TRUE |
| **PSD** | delta | 1 | 4 | 1 | concentrative | 5 | PRE | POST | 3.97557802270321 | 4.71076676368434 | paired_ttest | -1.26216728331798 | 4 | 0.56445836889505 | 0.275465345989441 | 0.275465345989441 | FALSE |
| **PSD** | delta | 1 | 4 | 1 | concentrative | 5 | ON | POST | 12.4489794168014 | 4.71076676368434 | paired_ttest | 18.5995672925299 | 4 | -8.3179793636357 | 4.91831842587904e-05 | 0.000147549552776371 | TRUE |
| **PSD** | delta | 1 | 4 | 2 | nap | 9 | PRE | ON | 2.37268408965744 | 11.7583436465878 | paired_ttest | -12.6673902599394 | 8 | 4.22246341997979 | 1.41782640586084e-06 | 4.25347921758252e-06 | TRUE |
| **PSD** | delta | 1 | 4 | 2 | nap | 9 | PRE | POST | 2.37268408965744 | 3.17068147701621 | paired_ttest | -0.56169119418806 | 8 | 0.187230398062687 | 0.589711306914455 | 0.589711306914455 | FALSE |
| **PSD** | delta | 1 | 4 | 2 | nap | 9 | ON | POST | 11.7583436465878 | 3.17068147701621 | paired_ttest | 7.5458555470234 | 8 | -2.51528518234113 | 6.63406964175984e-05 | 0.000132681392835197 | TRUE |
| **PSD** | delta | 1 | 4 | 3 | reading | 4 | PRE | ON | 1.40207301365496 | 6.87961214001714 | paired_ttest | -3.09504491830623 | 3 | 1.54752245915311 | 0.0535022117572664 | 0.107004423514533 | FALSE |
| **PSD** | delta | 1 | 4 | 3 | reading | 4 | PRE | POST | 1.40207301365496 | 0.736066205028252 | paired_ttest | 0.367167293823523 | 3 | -0.183583646911761 | 0.737865713225201 | 0.737865713225201 | FALSE |
| **PSD** | delta | 1 | 4 | 3 | reading | 4 | ON | POST | 6.87961214001714 | 0.736066205028252 | paired_ttest | 11.4081291455035 | 3 | -5.70406457275173 | 0.00144524977310362 | 0.00433574931931086 | TRUE |
| **PSD** | delta | 1 | 4 | 4 | analytical | 5 | PRE | ON | 2.31232535164667 | 9.70634178643025 | paired_ttest | -7.57064871512253 | 4 | 3.38569703215708 | 0.001631995973527 | 0.003263991947054 | TRUE |
| **PSD** | delta | 1 | 4 | 4 | analytical | 5 | PRE | POST | 2.31232535164667 | 1.23203540743099 | paired_ttest | 1.26252427030837 | 4 | -0.564618018330567 | 0.275349561268167 | 0.275349561268167 | FALSE |
| **PSD** | delta | 1 | 4 | 4 | analytical | 5 | ON | POST | 9.70634178643025 | 1.23203540743099 | paired_ttest | 32.2022539683641 | 4 | -14.4012857803949 | 5.54394970771014e-06 | 1.66318491231304e-05 | TRUE |
| **PSD** | delta | 1 | 4 | 5 | diss. el. 1 | 16 | PRE | ON | 2.00250085610972 | 10.3201184481551 | paired_ttest | -10.9954654047146 | 15 | 2.74886635117866 | 1.41406680058769e-08 | 2.82813360117537e-08 | TRUE |
| **PSD** | delta | 1 | 4 | 5 | diss. el. 1 | 16 | PRE | POST | 2.00250085610972 | 1.33655105260326 | paired_ttest | 0.685030871966157 | 15 | -0.171257717991539 | 0.503773437893268 | 0.503773437893268 | FALSE |
| **PSD** | delta | 1 | 4 | 5 | diss. el. 1 | 16 | ON | POST | 10.3201184481551 | 1.33655105260326 | paired_ttest | 14.2553248392337 | 15 | -3.56383120980843 | 3.9748292508833e-10 | 1.19244877526499e-09 | TRUE |
| **PSD** | delta | 1 | 4 | 6 | diss. el. 2 | 20 | PRE | ON | 5.37547725338474 | 12.5733176122033 | paired_ttest | -44.0667051701133 | 19 | 9.85361483048147 | 1.34477548554114e-20 | 4.03432645662342e-20 | TRUE |
| **PSD** | delta | 1 | 4 | 6 | diss. el. 2 | 20 | PRE | POST | 5.37547725338474 | 4.28190074672309 | paired_ttest | 6.29722659864232 | 19 | -1.4081026744284 | 4.80471306539047e-06 | 4.80471306539047e-06 | TRUE |
| **PSD** | delta | 1 | 4 | 6 | diss. el. 2 | 20 | ON | POST | 12.5733176122033 | 4.28190074672309 | paired_ttest | 36.1784599846949 | 19 | -8.08974958470338 | 5.46800075986147e-19 | 1.09360015197229e-18 | TRUE |
| **PSD** | delta | 1 | 4 | 7 | diss. el. 3 | 24 | PRE | ON | 3.1136406873458 | 10.3388685169166 | paired_ttest | -35.2837293040657 | 23 | 7.20226108478726 | 1.56401416503957e-21 | 3.12802833007915e-21 | TRUE |
| **PSD** | delta | 1 | 4 | 7 | diss. el. 3 | 24 | PRE | POST | 3.1136406873458 | 1.59369396336121 | paired_ttest | 7.08999421267152 | 23 | -1.44723900836091 | 3.19067313543465e-07 | 3.19067313543465e-07 | TRUE |
| **PSD** | delta | 1 | 4 | 7 | diss. el. 3 | 24 | ON | POST | 10.3388685169166 | 1.59369396336121 | paired_ttest | 37.9563984232144 | 23 | -7.74781738422127 | 2.99730539049409e-22 | 8.99191617148226e-22 | TRUE |
| **PSD** | gamma | 30 | 45 | 1 | concentrative | 5 | PRE | ON | -9.05386196943989 | -7.81769425333846 | paired_ttest | -3.91674586812967 | 4 | 1.75162200234587 | 0.0172950818199229 | 0.0518852454597686 | FALSE |
| **PSD** | gamma | 30 | 45 | 1 | concentrative | 5 | PRE | POST | -9.05386196943989 | -8.4542443541533 | paired_ttest | -0.62959807476859 | 4 | 0.281564818737112 | 0.56312850373243 | 0.875017308147925 | FALSE |
| **PSD** | gamma | 30 | 45 | 1 | concentrative | 5 | ON | POST | -7.81769425333846 | -8.4542443541533 | paired_ttest | 0.86157250919528 | 4 | -0.385306939621142 | 0.437508654073962 | 0.875017308147925 | FALSE |
| **PSD** | gamma | 30 | 45 | 2 | nap | 9 | PRE | ON | -12.86541292674 | -12.146815777991 | paired_ttest | -1.53516887380599 | 8 | 0.511722957935331 | 0.16329071791791 | 0.489872153753731 | FALSE |
| **PSD** | gamma | 30 | 45 | 2 | nap | 9 | PRE | POST | -12.86541292674 | -12.2934827044876 | paired_ttest | -0.421279001564875 | 8 | 0.140426333854958 | 0.684642083287613 | 1 | FALSE |
| **PSD** | gamma | 30 | 45 | 2 | nap | 9 | ON | POST | -12.146815777991 | -12.2934827044876 | paired_ttest | 0.0996718598460761 | 8 | -0.0332239532820254 | 0.923057274395085 | 1 | FALSE |
| **PSD** | gamma | 30 | 45 | 3 | reading | 4 | PRE | ON | -9.86263682990617 | -9.50501142302668 | paired_ttest | -0.679733877030897 | 3 | 0.339866938515448 | 0.54542443059916 | 1 | FALSE |
| **PSD** | gamma | 30 | 45 | 3 | reading | 4 | PRE | POST | -9.86263682990617 | -10.0335787999546 | paired_ttest | 0.133910442206881 | 3 | -0.0669552211034406 | 0.901951905804876 | 1 | FALSE |
| **PSD** | gamma | 30 | 45 | 3 | reading | 4 | ON | POST | -9.50501142302668 | -10.0335787999546 | paired_ttest | 0.615634104592047 | 3 | -0.307817052296024 | 0.581690600105163 | 1 | FALSE |
| **PSD** | gamma | 30 | 45 | 4 | analytical | 5 | PRE | ON | -11.7798187158681 | -11.1035352073672 | paired_ttest | -1.27526473640451 | 4 | 0.570315727981769 | 0.271247765834408 | 0.813743297503223 | FALSE |
| **PSD** | gamma | 30 | 45 | 4 | analytical | 5 | PRE | POST | -11.7798187158681 | -10.7757998199186 | paired_ttest | -1.18432770174419 | 4 | 0.529647449747223 | 0.301848040153718 | 0.813743297503223 | FALSE |
| **PSD** | gamma | 30 | 45 | 4 | analytical | 5 | ON | POST | -11.1035352073672 | -10.7757998199186 | paired_ttest | -0.579149927001491 | 4 | 0.259003721187875 | 0.593537868557591 | 0.813743297503223 | FALSE |
| **PSD** | gamma | 30 | 45 | 5 | diss. el. 1 | 16 | PRE | ON | -10.3477062334896 | -10.7943100484392 | paired_ttest | 0.67540874134364 | 15 | -0.16885218533591 | 0.509694849661319 | 0.656682196323069 | FALSE |
| **PSD** | gamma | 30 | 45 | 5 | diss. el. 1 | 16 | PRE | POST | -10.3477062334896 | -11.2133243175112 | paired_ttest | 1.01036666743253 | 15 | -0.252591666858133 | 0.328341098161535 | 0.656682196323069 | FALSE |
| **PSD** | gamma | 30 | 45 | 5 | diss. el. 1 | 16 | ON | POST | -10.7943100484392 | -11.2133243175112 | paired_ttest | 1.29876249781299 | 15 | -0.324690624453247 | 0.213637003813824 | 0.640911011441471 | FALSE |
| **PSD** | gamma | 30 | 45 | 6 | diss. el. 2 | 20 | PRE | ON | -8.76851341267933 | -8.34675722975457 | paired_ttest | -1.30666676666666 | 19 | 0.29217957142065 | 0.20692176085978 | 0.620765282579341 | FALSE |
| **PSD** | gamma | 30 | 45 | 6 | diss. el. 2 | 20 | PRE | POST | -8.76851341267933 | -8.01338082069822 | paired_ttest | -1.28837924960538 | 19 | 0.28809035829178 | 0.213083020564169 | 0.620765282579341 | FALSE |
| **PSD** | gamma | 30 | 45 | 6 | diss. el. 2 | 20 | ON | POST | -8.34675722975457 | -8.01338082069822 | paired_ttest | -0.602388261553557 | 19 | 0.134698110168168 | 0.554037702516432 | 0.620765282579341 | FALSE |
| **PSD** | gamma | 30 | 45 | 7 | diss. el. 3 | 24 | PRE | ON | -7.36847755054467 | -6.39750299975347 | paired_ttest | -2.5377677898333 | 23 | 0.51801968089685 | 0.0183915984381683 | 0.055174795314505 | FALSE |
| **PSD** | gamma | 30 | 45 | 7 | diss. el. 3 | 24 | PRE | POST | -7.36847755054467 | -7.12192837798594 | paired_ttest | -0.543586264212506 | 23 | 0.110959081542197 | 0.591956636252552 | 0.591956636252552 | FALSE |
| **PSD** | gamma | 30 | 45 | 7 | diss. el. 3 | 24 | ON | POST | -6.39750299975347 | -7.12192837798594 | paired_ttest | 1.63060061224784 | 23 | -0.332844956189755 | 0.116594211817028 | 0.233188423634057 | FALSE |
| **PSD** | theta | 4 | 8 | 1 | concentrative | 5 | PRE | ON | 0.479312479362295 | 13.1054397332959 | paired_ttest | -50.0836880161527 | 4 | 22.3981061936018 | 9.51070418183403e-07 | 1.90214083636681e-06 | TRUE |
| **PSD** | theta | 4 | 8 | 1 | concentrative | 5 | PRE | POST | 0.479312479362295 | 0.734327255998538 | paired_ttest | -0.654247700046279 | 4 | 0.292588466285274 | 0.548660370626199 | 0.548660370626199 | FALSE |
| **PSD** | theta | 4 | 8 | 1 | concentrative | 5 | ON | POST | 13.1054397332959 | 0.734327255998538 | paired_ttest | 56.9033930857187 | 4 | -25.4479710180117 | 5.71090533828033e-07 | 1.7132716014841e-06 | TRUE |
| **PSD** | theta | 4 | 8 | 2 | nap | 9 | PRE | ON | 0.211707227272442 | 13.8150014808412 | paired_ttest | -34.9929343705712 | 8 | 11.6643114568571 | 4.86623769382686e-10 | 1.45987130814806e-09 | TRUE |
| **PSD** | theta | 4 | 8 | 2 | nap | 9 | PRE | POST | 0.211707227272442 | 0.623610353644215 | paired_ttest | -0.464205645964284 | 8 | 0.154735215321428 | 0.654871027052854 | 0.654871027052854 | FALSE |
| **PSD** | theta | 4 | 8 | 2 | nap | 9 | ON | POST | 13.8150014808412 | 0.623610353644215 | paired_ttest | 19.4401648136625 | 8 | -6.48005493788751 | 5.09169141993042e-08 | 1.01833828398608e-07 | TRUE |
| **PSD** | theta | 4 | 8 | 3 | reading | 4 | PRE | ON | -1.34519040338251 | 8.56272535172265 | paired_ttest | -3.61532677914195 | 3 | 1.80766338957098 | 0.0363668588479797 | 0.0727337176959594 | FALSE |
| **PSD** | theta | 4 | 8 | 3 | reading | 4 | PRE | POST | -1.34519040338251 | -2.74495292625527 | paired_ttest | 0.423375805767601 | 3 | -0.2116879028838 | 0.700540420112768 | 0.700540420112768 | FALSE |
| **PSD** | theta | 4 | 8 | 3 | reading | 4 | ON | POST | 8.56272535172265 | -2.74495292625527 | paired_ttest | 10.9928266004101 | 3 | -5.49641330020504 | 0.00161195760654152 | 0.00483587281962457 | TRUE |
| **PSD** | theta | 4 | 8 | 4 | analytical | 5 | PRE | ON | 0.200142315711348 | 11.4733048383878 | paired_ttest | -9.38563980095117 | 4 | 4.19738572145088 | 0.000717997450474144 | 0.00143599490094829 | TRUE |
| **PSD** | theta | 4 | 8 | 4 | analytical | 5 | PRE | POST | 0.200142315711348 | -1.2369427018654 | paired_ttest | 1.24284009451151 | 4 | -0.555814987298001 | 0.281803875344568 | 0.281803875344568 | FALSE |
| **PSD** | theta | 4 | 8 | 4 | analytical | 5 | ON | POST | 11.4733048383878 | -1.2369427018654 | paired_ttest | 52.2183252993795 | 4 | -23.3527450081219 | 8.05004721435475e-07 | 2.41501416430642e-06 | TRUE |
| **PSD** | theta | 4 | 8 | 5 | diss. el. 1 | 16 | PRE | ON | 0.38128702297601 | 12.4529213624079 | paired_ttest | -13.7683179360278 | 15 | 3.44207948400695 | 6.46578166506301e-10 | 1.2931563330126e-09 | TRUE |
| **PSD** | theta | 4 | 8 | 5 | diss. el. 1 | 16 | PRE | POST | 0.38128702297601 | 0.204592000013842 | paired_ttest | 0.164069696087644 | 15 | -0.0410174240219111 | 0.87186662525934 | 0.87186662525934 | FALSE |
| **PSD** | theta | 4 | 8 | 5 | diss. el. 1 | 16 | ON | POST | 12.4529213624079 | 0.204592000013842 | paired_ttest | 16.9197889513344 | 15 | -4.22994723783361 | 3.50620044243172e-11 | 1.05186013272952e-10 | TRUE |
| **PSD** | theta | 4 | 8 | 6 | diss. el. 2 | 20 | PRE | ON | 2.73082858082078 | 13.5547622577357 | paired_ttest | -49.220931900699 | 19 | 11.0061349645851 | 1.67289185050115e-21 | 3.3457837010023e-21 | TRUE |
| **PSD** | theta | 4 | 8 | 6 | diss. el. 2 | 20 | PRE | POST | 2.73082858082078 | 2.0292159400072 | paired_ttest | 4.62611437631712 | 19 | -1.03443062171341 | 0.000184248943329852 | 0.000184248943329852 | TRUE |
| **PSD** | theta | 4 | 8 | 6 | diss. el. 2 | 20 | ON | POST | 13.5547622577357 | 2.0292159400072 | paired_ttest | 60.8270026426282 | 19 | -13.6013312776476 | 3.06939052302806e-23 | 9.20817156908418e-23 | TRUE |
| **PSD** | theta | 4 | 8 | 7 | diss. el. 3 | 24 | PRE | ON | 0.10559750212431 | 11.5017504977138 | paired_ttest | -64.2129880186424 | 23 | 13.1074212920936 | 1.88031434029076e-27 | 3.76062868058153e-27 | TRUE |
| **PSD** | theta | 4 | 8 | 7 | diss. el. 3 | 24 | PRE | POST | 0.10559750212431 | -0.678463053822607 | paired_ttest | 2.70206164737126 | 23 | -0.551556024133643 | 0.0127193061958002 | 0.0127193061958002 | TRUE |
| **PSD** | theta | 4 | 8 | 7 | diss. el. 3 | 24 | ON | POST | 11.5017504977138 | -0.678463053822607 | paired_ttest | 66.7846101333351 | 23 | -13.6323514581148 | 7.65612215195101e-28 | 2.2968366455853e-27 | TRUE |

**Supplementary Table S4. Repeated-measures ANOVA and post-hoc comparisons of global dDTF across PRE, ON and POST periods**

============================================================
GLOBAL CONNECTIVITY PRE / ON / POST TESTS ACROSS ALL BANDS AND SESSIONS
============================================================

| **Analysis** | **BandName** | **BandLowHz** | **BandHighHz** | **SessionIndex** | **SessionName** | **NPairedChunks** | **GlobalTest** | **Statistic** | **DF1** | **DF2** | **PValue** | **PAdjusted** | **Significant** |
| --- | --- | --- | --- | --- | --- | --- | --- | --- | --- | --- | --- | --- | --- |
| **Connectivity** | alpha | 8 | 13 | 1 | concentrative | 5 | rm_anova | 0.630747851591027 | 2 | 8 | 0.556718213276282 | 0.590458711050602 | FALSE |
| **Connectivity** | alpha | 8 | 13 | 2 | nap | 9 | rm_anova | 0.106807550367516 | 2 | 16 | 0.899333952185054 | 0.899333952185054 | FALSE |
| **Connectivity** | alpha | 8 | 13 | 3 | reading | 4 | rm_anova | 7.06163359516753 | 2 | 6 | 0.0265068591382115 | 0.0343607433273112 | TRUE |
| **Connectivity** | alpha | 8 | 13 | 4 | analytical | 5 | rm_anova | 0.668524845327288 | 2 | 8 | 0.538916235781708 | 0.589439632886243 | FALSE |
| **Connectivity** | alpha | 8 | 13 | 5 | diss. el. 1 | 16 | rm_anova | 0.263023002734705 | 2 | 30 | 0.770478465140071 | 0.79313959646772 | FALSE |
| **Connectivity** | alpha | 8 | 13 | 6 | diss. el. 2 | 20 | rm_anova | 6.23138827797584 | 2 | 38 | 0.0045649563338968 | 0.00694667268201687 | TRUE |
| **Connectivity** | alpha | 8 | 13 | 7 | diss. el. 3 | 24 | rm_anova | 9.19190648008477 | 2 | 46 | 0.000438092873561686 | 0.000851847254147723 | TRUE |
| **Connectivity** | beta | 13 | 30 | 1 | concentrative | 5 | rm_anova | 11.1131242288597 | 2 | 8 | 0.00490707794408028 | 0.00715615533511707 | TRUE |
| **Connectivity** | beta | 13 | 30 | 2 | nap | 9 | rm_anova | 20.8847788089811 | 2 | 16 | 3.46232256668555e-05 | 8.07875265559961e-05 | TRUE |
| **Connectivity** | beta | 13 | 30 | 3 | reading | 4 | rm_anova | 0.795327283772849 | 2 | 6 | 0.493873890025888 | 0.557599553255034 | FALSE |
| **Connectivity** | beta | 13 | 30 | 4 | analytical | 5 | rm_anova | 5.08306525891035 | 2 | 8 | 0.0376105970288487 | 0.0470132462860609 | TRUE |
| **Connectivity** | beta | 13 | 30 | 5 | diss. el. 1 | 16 | rm_anova | 27.3412544085098 | 2 | 30 | 1.73745306128243e-07 | 4.34363265320608e-07 | TRUE |
| **Connectivity** | beta | 13 | 30 | 6 | diss. el. 2 | 20 | rm_anova | 59.1736044209312 | 2 | 38 | 2.12900076946835e-12 | 1.06450038473418e-11 | TRUE |
| **Connectivity** | beta | 13 | 30 | 7 | diss. el. 3 | 24 | rm_anova | 76.4290404539173 | 2 | 46 | 2.38196578371827e-15 | 1.66737604860279e-14 | TRUE |
| **Connectivity** | delta | 1 | 4 | 1 | concentrative | 5 | rm_anova | 20.6011563505622 | 2 | 8 | 0.000698904570922173 | 0.00128745578854085 | TRUE |
| **Connectivity** | delta | 1 | 4 | 2 | nap | 9 | rm_anova | 15.0269014693427 | 2 | 16 | 0.000212244221323921 | 0.000436973396843367 | TRUE |
| **Connectivity** | delta | 1 | 4 | 3 | reading | 4 | rm_anova | 3.13324673289458 | 2 | 6 | 0.11702872018665 | 0.141241558845957 | FALSE |
| **Connectivity** | delta | 1 | 4 | 4 | analytical | 5 | rm_anova | 11.8743921411794 | 2 | 8 | 0.0040313596796896 | 0.00641352676314254 | TRUE |
| **Connectivity** | delta | 1 | 4 | 5 | diss. el. 1 | 16 | rm_anova | 36.6842079451319 | 2 | 30 | 8.72973194884803e-09 | 2.77764198372437e-08 | TRUE |
| **Connectivity** | delta | 1 | 4 | 6 | diss. el. 2 | 20 | rm_anova | 152.848731415281 | 2 | 38 | 6.73811064315949e-19 | 5.89584681276455e-18 | TRUE |
| **Connectivity** | delta | 1 | 4 | 7 | diss. el. 3 | 24 | rm_anova | 200.087350422126 | 2 | 46 | 2.01751365479098e-23 | 2.35376593058948e-22 | TRUE |
| **Connectivity** | gamma | 30 | 45 | 1 | concentrative | 5 | rm_anova | 12.8805505040989 | 2 | 8 | 0.00315277909956113 | 0.00525463183260189 | TRUE |
| **Connectivity** | gamma | 30 | 45 | 2 | nap | 9 | rm_anova | 9.90258659618298 | 2 | 16 | 0.00158998721055334 | 0.00278247761846835 | TRUE |
| **Connectivity** | gamma | 30 | 45 | 3 | reading | 4 | rm_anova | 2.99425780278636 | 2 | 6 | 0.125359575357059 | 0.146252837916569 | FALSE |
| **Connectivity** | gamma | 30 | 45 | 4 | analytical | 5 | rm_anova | 5.99909732853509 | 2 | 8 | 0.0256092454421057 | 0.0343607433273112 | TRUE |
| **Connectivity** | gamma | 30 | 45 | 5 | diss. el. 1 | 16 | rm_anova | 12.5032188096146 | 2 | 30 | 0.000112361111311212 | 0.000245789930993277 | TRUE |
| **Connectivity** | gamma | 30 | 45 | 6 | diss. el. 2 | 20 | rm_anova | 43.3433928919693 | 2 | 38 | 1.56783433614478e-10 | 6.09713352945193e-10 | TRUE |
| **Connectivity** | gamma | 30 | 45 | 7 | diss. el. 3 | 24 | rm_anova | 23.5422209875085 | 2 | 46 | 9.10451547616437e-08 | 2.65548368054794e-07 | TRUE |
| **Connectivity** | theta | 4 | 8 | 1 | concentrative | 5 | rm_anova | 457.420881604482 | 2 | 8 | 5.64743169384207e-09 | 1.97660109284473e-08 | TRUE |
| **Connectivity** | theta | 4 | 8 | 2 | nap | 9 | rm_anova | 151.042814578863 | 2 | 16 | 4.09833517326468e-11 | 1.7930216383033e-10 | TRUE |
| **Connectivity** | theta | 4 | 8 | 3 | reading | 4 | rm_anova | 7.0624182042272 | 2 | 6 | 0.0265006590678785 | 0.0343607433273112 | TRUE |
| **Connectivity** | theta | 4 | 8 | 4 | analytical | 5 | rm_anova | 219.344855511131 | 2 | 8 | 1.02881230866643e-07 | 2.76987929256347e-07 | TRUE |
| **Connectivity** | theta | 4 | 8 | 5 | diss. el. 1 | 16 | rm_anova | 113.74247431687 | 2 | 30 | 9.89785554963418e-15 | 5.77374907061994e-14 | TRUE |
| **Connectivity** | theta | 4 | 8 | 6 | diss. el. 2 | 20 | rm_anova | 880.604481180778 | 2 | 38 | 1.47685489227821e-32 | 2.58449606148688e-31 | TRUE |
| **Connectivity** | theta | 4 | 8 | 7 | diss. el. 3 | 24 | rm_anova | 559.045216739807 | 2 | 46 | 5.31784364302992e-33 | 1.86124527506047e-31 | TRUE |

============================================================
CONNECTIVITY POSTHOC TESTS ACROSS ALL BANDS AND SESSIONS
============================================================

| **Analysis** | **BandName** | **BandLowHz** | **BandHighHz** | **SessionIndex** | **SessionName** | **NPairedChunks** | **Condition1** | **Condition2** | **Mean1** | **Mean2** | **Test** | **TestStatistic** | **DF** | **CohensDz** | **PValue** | **PAdjusted** | **Significant** |
| --- | --- | --- | --- | --- | --- | --- | --- | --- | --- | --- | --- | --- | --- | --- | --- | --- | --- |
| **Connectivity** | alpha | 8 | 13 | 1 | concentrative | 5 | PRE | ON | 0.00449043670401052 | 0.00438540351379405 | paired_ttest | 0.382750318057776 | 4 | -0.171171145917371 | 0.721374326476402 | 1 | FALSE |
| **Connectivity** | alpha | 8 | 13 | 1 | concentrative | 5 | PRE | POST | 0.00449043670401052 | 0.00419481424191536 | paired_ttest | 1.35889494421986 | 4 | -0.607716293911277 | 0.245754760998346 | 0.737264282995038 | FALSE |
| **Connectivity** | alpha | 8 | 13 | 1 | concentrative | 5 | ON | POST | 0.00438540351379405 | 0.00419481424191536 | paired_ttest | 0.631788698482583 | 4 | -0.282544495444635 | 0.561832230105548 | 1 | FALSE |
| **Connectivity** | alpha | 8 | 13 | 2 | nap | 9 | PRE | ON | 0.00364902318797257 | 0.00363998324297874 | paired_ttest | 0.0257225471404941 | 8 | -0.00857418238016471 | 0.980108700077774 | 1 | FALSE |
| **Connectivity** | alpha | 8 | 13 | 2 | nap | 9 | PRE | POST | 0.00364902318797257 | 0.00377405685027027 | paired_ttest | -0.386372625971882 | 8 | 0.128790875323961 | 0.709294056722582 | 1 | FALSE |
| **Connectivity** | alpha | 8 | 13 | 2 | nap | 9 | ON | POST | 0.00363998324297874 | 0.00377405685027027 | paired_ttest | -0.454025163185354 | 8 | 0.151341721061785 | 0.661874982610571 | 1 | FALSE |
| **Connectivity** | alpha | 8 | 13 | 3 | reading | 4 | PRE | ON | 0.00290999925909878 | 0.00485593871387935 | paired_ttest | -2.8985351440265 | 3 | 1.44926757201325 | 0.0625767390492192 | 0.125153478098438 | FALSE |
| **Connectivity** | alpha | 8 | 13 | 3 | reading | 4 | PRE | POST | 0.00290999925909878 | 0.00252340417566856 | paired_ttest | 0.506586423991343 | 3 | -0.253293211995672 | 0.647330860624885 | 0.647330860624885 | FALSE |
| **Connectivity** | alpha | 8 | 13 | 3 | reading | 4 | ON | POST | 0.00485593871387935 | 0.00252340417566856 | paired_ttest | 4.29625072234573 | 3 | -2.14812536117286 | 0.0231940564841383 | 0.0695821694524149 | FALSE |
| **Connectivity** | alpha | 8 | 13 | 4 | analytical | 5 | PRE | ON | 0.00482337703879788 | 0.0043955463925131 | paired_ttest | 1.24653634347295 | 4 | -0.557468000085908 | 0.280580985931677 | 0.6849692944112 | FALSE |
| **Connectivity** | alpha | 8 | 13 | 4 | analytical | 5 | PRE | POST | 0.00482337703879788 | 0.00468873342713734 | paired_ttest | 0.259538168583421 | 4 | -0.116068997541666 | 0.80803057039515 | 0.80803057039515 | FALSE |
| **Connectivity** | alpha | 8 | 13 | 4 | analytical | 5 | ON | POST | 0.0043955463925131 | 0.00468873342713734 | paired_ttest | -1.42113660209198 | 4 | 0.635551609518147 | 0.228323098137067 | 0.6849692944112 | FALSE |
| **Connectivity** | alpha | 8 | 13 | 5 | diss. el. 1 | 16 | PRE | ON | 0.00332641659598871 | 0.00338909184095999 | paired_ttest | -0.321674554274694 | 15 | 0.0804186385686734 | 0.752138242871747 | 1 | FALSE |
| **Connectivity** | alpha | 8 | 13 | 5 | diss. el. 1 | 16 | PRE | POST | 0.00332641659598871 | 0.00324281003355678 | paired_ttest | 0.349321251685046 | 15 | -0.0873303129212614 | 0.731703754604302 | 1 | FALSE |
| **Connectivity** | alpha | 8 | 13 | 5 | diss. el. 1 | 16 | ON | POST | 0.00338909184095999 | 0.00324281003355678 | paired_ttest | 0.880200391265612 | 15 | -0.220050097816403 | 0.392634963841434 | 1 | FALSE |
| **Connectivity** | alpha | 8 | 13 | 6 | diss. el. 2 | 20 | PRE | ON | 0.00679935791966299 | 0.00564389300593041 | paired_ttest | 4.49358520951955 | 19 | -1.00479619911734 | 0.000248779919436292 | 0.000746339758308877 | TRUE |
| **Connectivity** | alpha | 8 | 13 | 6 | diss. el. 2 | 20 | PRE | POST | 0.00679935791966299 | 0.00600589529479696 | paired_ttest | 1.89630040603058 | 19 | -0.424025661364482 | 0.0732315721651192 | 0.146463144330238 | FALSE |
| **Connectivity** | alpha | 8 | 13 | 6 | diss. el. 2 | 20 | ON | POST | 0.00564389300593041 | 0.00600589529479696 | paired_ttest | -1.17376240116 | 19 | 0.262461251842715 | 0.254994105666342 | 0.254994105666342 | FALSE |
| **Connectivity** | alpha | 8 | 13 | 7 | diss. el. 3 | 24 | PRE | ON | 0.00487317647717142 | 0.004278668624937 | paired_ttest | 4.21897601364449 | 23 | -0.861194872539204 | 0.000326239063930983 | 0.000978717191792949 | TRUE |
| **Connectivity** | alpha | 8 | 13 | 7 | diss. el. 3 | 24 | PRE | POST | 0.00487317647717142 | 0.00488153237534503 | paired_ttest | -0.0450776338476122 | 23 | 0.00920143347822183 | 0.964434524850334 | 0.964434524850334 | FALSE |
| **Connectivity** | alpha | 8 | 13 | 7 | diss. el. 3 | 24 | ON | POST | 0.004278668624937 | 0.00488153237534503 | paired_ttest | -3.90933586263073 | 23 | 0.797989841384034 | 0.000704416856838435 | 0.00140883371367687 | TRUE |
| **Connectivity** | beta | 13 | 30 | 1 | concentrative | 5 | PRE | ON | 0.0018527250254922 | 0.00145388451533922 | paired_ttest | 9.79734564462649 | 4 | -4.38150617208927 | 0.000608332164941622 | 0.00182499649482487 | TRUE |
| **Connectivity** | beta | 13 | 30 | 1 | concentrative | 5 | PRE | POST | 0.0018527250254922 | 0.0016854110042511 | paired_ttest | 2.01936552247666 | 4 | -0.903087715935438 | 0.113579820212132 | 0.226943196908018 | FALSE |
| **Connectivity** | beta | 13 | 30 | 1 | concentrative | 5 | ON | POST | 0.00145388451533922 | 0.0016854110042511 | paired_ttest | -2.02020221935664 | 4 | 0.903461898155477 | 0.113471598454009 | 0.226943196908018 | FALSE |
| **Connectivity** | beta | 13 | 30 | 2 | nap | 9 | PRE | ON | 0.00172916197565463 | 0.00133719306448283 | paired_ttest | 6.82785027897081 | 8 | -2.27595009299027 | 0.000133969301106926 | 0.000401907903320778 | TRUE |
| **Connectivity** | beta | 13 | 30 | 2 | nap | 9 | PRE | POST | 0.00172916197565463 | 0.00170860266869861 | paired_ttest | 0.287195665158198 | 8 | -0.095731888386066 | 0.781261005463676 | 0.781261005463676 | FALSE |
| **Connectivity** | beta | 13 | 30 | 2 | nap | 9 | ON | POST | 0.00133719306448283 | 0.00170860266869861 | paired_ttest | -4.98027021349434 | 8 | 1.66009007116478 | 0.00107912707637663 | 0.00215825415275325 | TRUE |
| **Connectivity** | beta | 13 | 30 | 3 | reading | 4 | PRE | ON | 0.00150667111532549 | 0.00151024919673697 | paired_ttest | -0.0321978188616727 | 3 | 0.0160989094308364 | 0.976336667186777 | 0.976336667186777 | FALSE |
| **Connectivity** | beta | 13 | 30 | 3 | reading | 4 | PRE | POST | 0.00150667111532549 | 0.00164557249739739 | paired_ttest | -0.807728426782344 | 3 | 0.403864213391172 | 0.478353009547536 | 0.956706019095071 | FALSE |
| **Connectivity** | beta | 13 | 30 | 3 | reading | 4 | ON | POST | 0.00151024919673697 | 0.00164557249739739 | paired_ttest | -1.84524531006145 | 3 | 0.922622655030723 | 0.162189383311986 | 0.486568149935957 | FALSE |
| **Connectivity** | beta | 13 | 30 | 4 | analytical | 5 | PRE | ON | 0.0016164033024101 | 0.00134234457771932 | paired_ttest | 5.69467491700715 | 4 | -2.54673604483819 | 0.00469766219620324 | 0.0140929865886097 | TRUE |
| **Connectivity** | beta | 13 | 30 | 4 | analytical | 5 | PRE | POST | 0.0016164033024101 | 0.00164733277355156 | paired_ttest | -0.25747127486089 | 4 | 0.115144654568496 | 0.809517843329742 | 0.809517843329742 | FALSE |
| **Connectivity** | beta | 13 | 30 | 4 | analytical | 5 | ON | POST | 0.00134234457771932 | 0.00164733277355156 | paired_ttest | -2.37309693581737 | 4 | 1.06128121313682 | 0.0765622775373212 | 0.153124555074642 | FALSE |
| **Connectivity** | beta | 13 | 30 | 5 | diss. el. 1 | 16 | PRE | ON | 0.00125956402501763 | 0.00101089708695217 | paired_ttest | 7.76294283318171 | 15 | -1.94073570829543 | 1.24471898043887e-06 | 3.73415694131662e-06 | TRUE |
| **Connectivity** | beta | 13 | 30 | 5 | diss. el. 1 | 16 | PRE | POST | 0.00125956402501763 | 0.00126079402948129 | paired_ttest | -0.0258984230483015 | 15 | 0.00647460576207538 | 0.979679766601852 | 0.979679766601852 | FALSE |
| **Connectivity** | beta | 13 | 30 | 5 | diss. el. 1 | 16 | ON | POST | 0.00101089708695217 | 0.00126079402948129 | paired_ttest | -7.02891568833961 | 15 | 1.7572289220849 | 4.07865176525181e-06 | 8.15730353050363e-06 | TRUE |
| **Connectivity** | beta | 13 | 30 | 6 | diss. el. 2 | 20 | PRE | ON | 0.00210866208459913 | 0.00168182594698054 | paired_ttest | 13.7572122878928 | 19 | -3.07620618566238 | 2.49434451928273e-11 | 7.48303355784819e-11 | TRUE |
| **Connectivity** | beta | 13 | 30 | 6 | diss. el. 2 | 20 | PRE | POST | 0.00210866208459913 | 0.00206981652385572 | paired_ttest | 0.851282635267214 | 19 | -0.190352584052265 | 0.4052141055653 | 0.4052141055653 | FALSE |
| **Connectivity** | beta | 13 | 30 | 6 | diss. el. 2 | 20 | ON | POST | 0.00168182594698054 | 0.00206981652385572 | paired_ttest | -7.60403441993856 | 19 | 1.70031378662308 | 3.53230537734599e-07 | 7.06461075469198e-07 | TRUE |
| **Connectivity** | beta | 13 | 30 | 7 | diss. el. 3 | 24 | PRE | ON | 0.00188601492534257 | 0.00155124390658707 | paired_ttest | 11.5225294409432 | 23 | -2.3520264730423 | 4.9480540987665e-11 | 1.48441622962995e-10 | TRUE |
| **Connectivity** | beta | 13 | 30 | 7 | diss. el. 3 | 24 | PRE | POST | 0.00188601492534257 | 0.00188420439701496 | paired_ttest | 0.0630704210250364 | 23 | -0.0128741957811536 | 0.950255355102734 | 0.950255355102734 | FALSE |
| **Connectivity** | beta | 13 | 30 | 7 | diss. el. 3 | 24 | ON | POST | 0.00155124390658707 | 0.00188420439701496 | paired_ttest | -9.42225697660917 | 23 | 1.92331015150595 | 2.32120922960447e-09 | 4.64241845920893e-09 | TRUE |
| **Connectivity** | delta | 1 | 4 | 1 | concentrative | 5 | PRE | ON | 0.0053209849995649 | 0.0069405377407355 | paired_ttest | -4.67038025963698 | 4 | 2.08865754826428 | 0.00951562034928107 | 0.0190312406985621 | TRUE |
| **Connectivity** | delta | 1 | 4 | 1 | concentrative | 5 | PRE | POST | 0.0053209849995649 | 0.00532327245638666 | paired_ttest | -0.00693939593956736 | 4 | 0.00310339220873173 | 0.994795505258432 | 0.994795505258432 | FALSE |
| **Connectivity** | delta | 1 | 4 | 1 | concentrative | 5 | ON | POST | 0.0069405377407355 | 0.00532327245638666 | paired_ttest | 10.1528947266999 | 4 | -4.54051255546003 | 0.000529923978613004 | 0.00158977193583901 | TRUE |
| **Connectivity** | delta | 1 | 4 | 2 | nap | 9 | PRE | ON | 0.00444965815685704 | 0.00624492672903906 | paired_ttest | -4.75420870789791 | 8 | 1.58473623596597 | 0.00143745389376123 | 0.00287490778752245 | TRUE |
| **Connectivity** | delta | 1 | 4 | 2 | nap | 9 | PRE | POST | 0.00444965815685704 | 0.00438040177688097 | paired_ttest | 0.161556216538614 | 8 | -0.0538520721795381 | 0.875660933629533 | 0.875660933629533 | FALSE |
| **Connectivity** | delta | 1 | 4 | 2 | nap | 9 | ON | POST | 0.00624492672903906 | 0.00438040177688097 | paired_ttest | 5.38723080080919 | 8 | -1.79574360026973 | 0.000655942775908606 | 0.00196782832772582 | TRUE |
| **Connectivity** | delta | 1 | 4 | 3 | reading | 4 | PRE | ON | 0.00636247413969957 | 0.00942123661413794 | paired_ttest | -1.99534136791207 | 3 | 0.997670683956033 | 0.139956651522875 | 0.27991330304575 | FALSE |
| **Connectivity** | delta | 1 | 4 | 3 | reading | 4 | PRE | POST | 0.00636247413969957 | 0.00615906940194063 | paired_ttest | 0.125690680333885 | 3 | -0.0628453401669423 | 0.90792697250874 | 0.90792697250874 | FALSE |
| **Connectivity** | delta | 1 | 4 | 3 | reading | 4 | ON | POST | 0.00942123661413794 | 0.00615906940194063 | paired_ttest | 2.73099780651937 | 3 | -1.36549890325968 | 0.0718803232164902 | 0.21564096964947 | FALSE |
| **Connectivity** | delta | 1 | 4 | 4 | analytical | 5 | PRE | ON | 0.00417106570256487 | 0.00566657397685924 | paired_ttest | -5.12158926645827 | 4 | 2.2904443505268 | 0.00687853654554894 | 0.0206356096366468 | TRUE |
| **Connectivity** | delta | 1 | 4 | 4 | analytical | 5 | PRE | POST | 0.00417106570256487 | 0.00404126393446647 | paired_ttest | 0.300205397797168 | 4 | -0.134255935337367 | 0.778975720614569 | 0.778975720614569 | FALSE |
| **Connectivity** | delta | 1 | 4 | 4 | analytical | 5 | ON | POST | 0.00566657397685924 | 0.00404126393446647 | paired_ttest | 4.34351521737381 | 4 | -1.94247905747052 | 0.0122191413519027 | 0.0244382827038053 | TRUE |
| **Connectivity** | delta | 1 | 4 | 5 | diss. el. 1 | 16 | PRE | ON | 0.00318565177803854 | 0.00487990618108074 | paired_ttest | -6.91277060518303 | 15 | 1.72819265129576 | 4.95356614450537e-06 | 9.90713228901074e-06 | TRUE |
| **Connectivity** | delta | 1 | 4 | 5 | diss. el. 1 | 16 | PRE | POST | 0.00318565177803854 | 0.0031379556257279 | paired_ttest | 0.162513033131383 | 15 | -0.0406282582828459 | 0.873070915454977 | 0.873070915454977 | FALSE |
| **Connectivity** | delta | 1 | 4 | 5 | diss. el. 1 | 16 | ON | POST | 0.00487990618108074 | 0.0031379556257279 | paired_ttest | 14.3086017676549 | 15 | -3.57715044191373 | 3.7721134442271e-10 | 1.13163403326813e-09 | TRUE |
| **Connectivity** | delta | 1 | 4 | 6 | diss. el. 2 | 20 | PRE | ON | 0.00504785551586522 | 0.00744035053249509 | paired_ttest | -15.2226667440227 | 19 | 3.40389176384601 | 4.24608512524904e-12 | 1.27382553757471e-11 | TRUE |
| **Connectivity** | delta | 1 | 4 | 6 | diss. el. 2 | 20 | PRE | POST | 0.00504785551586522 | 0.004978147557066 | paired_ttest | 0.57898065462135 | 19 | -0.129464010139067 | 0.569403337330223 | 0.569403337330223 | FALSE |
| **Connectivity** | delta | 1 | 4 | 6 | diss. el. 2 | 20 | ON | POST | 0.00744035053249509 | 0.004978147557066 | paired_ttest | 12.6396754756949 | 19 | -2.82631735771909 | 1.07246252713947e-10 | 2.14492505427894e-10 | TRUE |
| **Connectivity** | delta | 1 | 4 | 7 | diss. el. 3 | 24 | PRE | ON | 0.00445889741897749 | 0.00591367087434849 | paired_ttest | -15.959744771356 | 23 | 3.25776925957284 | 6.19554032303997e-14 | 1.23910806460799e-13 | TRUE |
| **Connectivity** | delta | 1 | 4 | 7 | diss. el. 3 | 24 | PRE | POST | 0.00445889741897749 | 0.00406861668676773 | paired_ttest | 3.56005101990329 | 23 | -0.726692371419825 | 0.0016651481210838 | 0.0016651481210838 | TRUE |
| **Connectivity** | delta | 1 | 4 | 7 | diss. el. 3 | 24 | ON | POST | 0.00591367087434849 | 0.00406861668676773 | paired_ttest | 20.5960601735216 | 23 | -4.20415317806553 | 2.54689237974256e-16 | 7.64067713922768e-16 | TRUE |
| **Connectivity** | gamma | 30 | 45 | 1 | concentrative | 5 | PRE | ON | 0.000711000978250922 | 0.000518121592261072 | paired_ttest | 4.63138185084824 | 4 | -2.07121692965109 | 0.00979705886872862 | 0.0293911766061859 | TRUE |
| **Connectivity** | gamma | 30 | 45 | 1 | concentrative | 5 | PRE | POST | 0.000711000978250922 | 0.000698554001601802 | paired_ttest | 0.271251065689953 | 4 | -0.121307164370399 | 0.799621054112068 | 0.799621054112068 | FALSE |
| **Connectivity** | gamma | 30 | 45 | 1 | concentrative | 5 | ON | POST | 0.000518121592261072 | 0.000698554001601802 | paired_ttest | -4.52815474097392 | 4 | 2.02505236269113 | 0.0105922307517263 | 0.0293911766061859 | TRUE |
| **Connectivity** | gamma | 30 | 45 | 2 | nap | 9 | PRE | ON | 0.000651401644599481 | 0.000386889914865712 | paired_ttest | 6.74684436159046 | 8 | -2.24894812053015 | 0.000145523925631079 | 0.000436571776893236 | TRUE |
| **Connectivity** | gamma | 30 | 45 | 2 | nap | 9 | PRE | POST | 0.000651401644599481 | 0.00061077000299363 | paired_ttest | 0.501666878800909 | 8 | -0.167222292933636 | 0.62941413036503 | 0.62941413036503 | FALSE |
| **Connectivity** | gamma | 30 | 45 | 2 | nap | 9 | ON | POST | 0.000386889914865712 | 0.00061077000299363 | paired_ttest | -3.45587653120214 | 8 | 1.15195884373405 | 0.00861992092733381 | 0.0172398418546676 | TRUE |
| **Connectivity** | gamma | 30 | 45 | 3 | reading | 4 | PRE | ON | 0.00107190331753001 | 0.000796377859816111 | paired_ttest | 2.78175609508873 | 3 | -1.39087804754436 | 0.0688898433589916 | 0.206669530076975 | FALSE |
| **Connectivity** | gamma | 30 | 45 | 3 | reading | 4 | PRE | POST | 0.00107190331753001 | 0.00128290328714081 | paired_ttest | -0.794487167998063 | 3 | 0.397243583999031 | 0.484958355412292 | 0.484958355412292 | FALSE |
| **Connectivity** | gamma | 30 | 45 | 3 | reading | 4 | ON | POST | 0.000796377859816111 | 0.00128290328714081 | paired_ttest | -2.46574982132948 | 3 | 1.23287491066474 | 0.0904046940204094 | 0.206669530076975 | FALSE |
| **Connectivity** | gamma | 30 | 45 | 4 | analytical | 5 | PRE | ON | 0.000668370255017416 | 0.000490569507591961 | paired_ttest | 2.44856357362481 | 4 | -1.09503091957098 | 0.0705543280896849 | 0.14110865617937 | FALSE |
| **Connectivity** | gamma | 30 | 45 | 4 | analytical | 5 | PRE | POST | 0.000668370255017416 | 0.000820022958904985 | paired_ttest | -1.25772758907144 | 4 | 0.56247287726813 | 0.276909212642276 | 0.276909212642276 | FALSE |
| **Connectivity** | gamma | 30 | 45 | 4 | analytical | 5 | ON | POST | 0.000490569507591961 | 0.000820022958904985 | paired_ttest | -3.83386899580394 | 4 | 1.71455833828929 | 0.0185565607421124 | 0.0556696822263372 | FALSE |
| **Connectivity** | gamma | 30 | 45 | 5 | diss. el. 1 | 16 | PRE | ON | 0.000752996235184965 | 0.000398885856627946 | paired_ttest | 4.66242669936638 | 15 | -1.1656066748416 | 0.000306645341974118 | 0.000613290683948236 | TRUE |
| **Connectivity** | gamma | 30 | 45 | 5 | diss. el. 1 | 16 | PRE | POST | 0.000752996235184965 | 0.000670635230919349 | paired_ttest | 0.87404018695362 | 15 | -0.218510046738405 | 0.395874808003842 | 0.395874808003842 | FALSE |
| **Connectivity** | gamma | 30 | 45 | 5 | diss. el. 1 | 16 | ON | POST | 0.000398885856627946 | 0.000670635230919349 | paired_ttest | -6.35236002951359 | 15 | 1.5880900073784 | 1.29829626082444e-05 | 3.89488878247332e-05 | TRUE |
| **Connectivity** | gamma | 30 | 45 | 6 | diss. el. 2 | 20 | PRE | ON | 0.000726020919574226 | 0.000506082584596481 | paired_ttest | 10.4229874828958 | 19 | -2.33065085403844 | 2.68922099110394e-09 | 8.06766297331182e-09 | TRUE |
| **Connectivity** | gamma | 30 | 45 | 6 | diss. el. 2 | 20 | PRE | POST | 0.000726020919574226 | 0.000821460372710814 | paired_ttest | -2.15895293112003 | 19 | 0.48275655142068 | 0.0438522732166689 | 0.0438522732166689 | TRUE |
| **Connectivity** | gamma | 30 | 45 | 6 | diss. el. 2 | 20 | ON | POST | 0.000506082584596481 | 0.000821460372710814 | paired_ttest | -9.02371561936024 | 19 | 2.01776415345161 | 2.6806678948287e-08 | 5.3613357896574e-08 | TRUE |
| **Connectivity** | gamma | 30 | 45 | 7 | diss. el. 3 | 24 | PRE | ON | 0.000924864401063906 | 0.000664088952593454 | paired_ttest | 5.95614047810454 | 23 | -1.2157920839744 | 4.50691518497779e-06 | 9.01383036995557e-06 | TRUE |
| **Connectivity** | gamma | 30 | 45 | 7 | diss. el. 3 | 24 | PRE | POST | 0.000924864401063906 | 0.00099697131801168 | paired_ttest | -1.26273248994306 | 23 | 0.257754190166216 | 0.219338030533599 | 0.219338030533599 | FALSE |
| **Connectivity** | gamma | 30 | 45 | 7 | diss. el. 3 | 24 | ON | POST | 0.000664088952593454 | 0.00099697131801168 | paired_ttest | -6.48047496231592 | 23 | 1.32282141237967 | 1.29941712145293e-06 | 3.89825136435878e-06 | TRUE |
| **Connectivity** | theta | 4 | 8 | 1 | concentrative | 5 | PRE | ON | 0.00236075143022901 | 0.00659594966013828 | paired_ttest | -19.7952619786937 | 4 | 8.85271028335523 | 3.84196490716875e-05 | 7.6839298143375e-05 | TRUE |
| **Connectivity** | theta | 4 | 8 | 1 | concentrative | 5 | PRE | POST | 0.00236075143022901 | 0.00231750936249457 | paired_ttest | 0.294228263168104 | 4 | -0.131582879469116 | 0.783220319601847 | 0.783220319601847 | FALSE |
| **Connectivity** | theta | 4 | 8 | 1 | concentrative | 5 | ON | POST | 0.00659594966013828 | 0.00231750936249457 | paired_ttest | 39.2848033998966 | 4 | -17.5686981769767 | 2.50829834938717e-06 | 7.52489504816152e-06 | TRUE |
| **Connectivity** | theta | 4 | 8 | 2 | nap | 9 | PRE | ON | 0.00283519195821203 | 0.0073830195445417 | paired_ttest | -13.3571227615011 | 8 | 4.45237425383371 | 9.43866274534793e-07 | 2.83159882360438e-06 | TRUE |
| **Connectivity** | theta | 4 | 8 | 2 | nap | 9 | PRE | POST | 0.00283519195821203 | 0.00314428505517646 | paired_ttest | -2.28778742525186 | 8 | 0.762595808417287 | 0.0514422109682442 | 0.0514422109682442 | FALSE |
| **Connectivity** | theta | 4 | 8 | 2 | nap | 9 | ON | POST | 0.0073830195445417 | 0.00314428505517646 | paired_ttest | 12.1186989992164 | 8 | -4.03956633307215 | 1.9888686468585e-06 | 3.97773729371699e-06 | TRUE |
| **Connectivity** | theta | 4 | 8 | 3 | reading | 4 | PRE | ON | 0.00403484230716468 | 0.0105813639098371 | paired_ttest | -3.5705092570045 | 3 | 1.78525462850225 | 0.037538477192147 | 0.0768858760957299 | FALSE |
| **Connectivity** | theta | 4 | 8 | 3 | reading | 4 | PRE | POST | 0.00403484230716468 | 0.00367497905118104 | paired_ttest | 0.139106359483782 | 3 | -0.069553179741891 | 0.898179377296325 | 0.898179377296325 | FALSE |
| **Connectivity** | theta | 4 | 8 | 3 | reading | 4 | ON | POST | 0.0105813639098371 | 0.00367497905118104 | paired_ttest | 4.13745619292792 | 3 | -2.06872809646396 | 0.0256286253652433 | 0.0768858760957299 | FALSE |
| **Connectivity** | theta | 4 | 8 | 4 | analytical | 5 | PRE | ON | 0.00236594272679765 | 0.00657223306340665 | paired_ttest | -16.2924608555064 | 4 | 7.28620999873335 | 8.30567885168513e-05 | 0.000218761355770595 | TRUE |
| **Connectivity** | theta | 4 | 8 | 4 | analytical | 5 | PRE | POST | 0.00236594272679765 | 0.00233038477696084 | paired_ttest | 0.196992629020265 | 4 | -0.0880977819111419 | 0.853437941415768 | 0.853437941415768 | FALSE |
| **Connectivity** | theta | 4 | 8 | 4 | analytical | 5 | ON | POST | 0.00657223306340665 | 0.00233038477696084 | paired_ttest | 16.8379632736057 | 4 | -7.53016609648544 | 7.29204519235318e-05 | 0.000218761355770595 | TRUE |
| **Connectivity** | theta | 4 | 8 | 5 | diss. el. 1 | 16 | PRE | ON | 0.00192893978640587 | 0.00571449216810268 | paired_ttest | -11.5629586530633 | 15 | 2.89073966326581 | 7.16528166655732e-09 | 1.43305633331146e-08 | TRUE |
| **Connectivity** | theta | 4 | 8 | 5 | diss. el. 1 | 16 | PRE | POST | 0.00192893978640587 | 0.00202400583048381 | paired_ttest | -0.278857789875205 | 15 | 0.0697144474688011 | 0.7841617561199 | 0.7841617561199 | FALSE |
| **Connectivity** | theta | 4 | 8 | 5 | diss. el. 1 | 16 | ON | POST | 0.00571449216810268 | 0.00202400583048381 | paired_ttest | 24.6537071815652 | 15 | -6.1634267953913 | 1.49406421907022e-13 | 4.48219265721065e-13 | TRUE |
| **Connectivity** | theta | 4 | 8 | 6 | diss. el. 2 | 20 | PRE | ON | 0.00299566957195131 | 0.00801126510728802 | paired_ttest | -30.6945335531222 | 19 | 6.86350635624295 | 1.18139495110278e-17 | 2.36278990220556e-17 | TRUE |
| **Connectivity** | theta | 4 | 8 | 6 | diss. el. 2 | 20 | PRE | POST | 0.00299566957195131 | 0.00300224000375548 | paired_ttest | -0.0670396772419804 | 19 | 0.0149905275502714 | 0.94725055364155 | 0.94725055364155 | FALSE |
| **Connectivity** | theta | 4 | 8 | 6 | diss. el. 2 | 20 | ON | POST | 0.00801126510728802 | 0.00300224000375548 | paired_ttest | 34.7707482425712 | 19 | -7.77497566989206 | 1.14992944815652e-18 | 3.44978834446957e-18 | TRUE |
| **Connectivity** | theta | 4 | 8 | 7 | diss. el. 3 | 24 | PRE | ON | 0.00259545408443535 | 0.0058488042929352 | paired_ttest | -27.5352574342177 | 23 | 5.62061088750088 | 4.12559499717569e-19 | 9.46490051468295e-19 | TRUE |
| **Connectivity** | theta | 4 | 8 | 7 | diss. el. 3 | 24 | PRE | POST | 0.00259545408443535 | 0.00254432313219627 | paired_ttest | 0.500193627918821 | 23 | -0.102101596749388 | 0.621691052603559 | 0.621691052603559 | FALSE |
| **Connectivity** | theta | 4 | 8 | 7 | diss. el. 3 | 24 | ON | POST | 0.0058488042929352 | 0.00254432313219627 | paired_ttest | 27.8676009748957 | 23 | -5.68845022866514 | 3.15496683822765e-19 | 9.46490051468295e-19 | TRUE |

**S5. Transcription of volunteer’s interview. Selected Excerpts.**

This interview forms part of the qualitative material accompanying the present study on advanced Vajrayāna meditation. The interview was conducted by one of the authors in English. The volunteer replied in Tibetan, and consecutive interpretation between English and Tibetan was provided by interpreters from Sera Jey Monastic University. The text below is an editorially revised English version intended for scientific publication. Time stamps refer to the original audiovisual recording.

**00:00:00–00:00:38**

**Interviewer:** Thank you for agreeing to participate. Your experience is extremely valuable for our research. I will begin with a few questions about your background and monastic education, and then we will focus on your meditation practice, particularly the Dissolution of the Elements. Is that acceptable to you?

[Sections unrelated to the present study have been omitted.]

**00:09:12–00:09:32**

**Interviewer:** In Vajrayāna there are meditation practices that, as I understand it, are intended to prepare practitioners for the process of dying. One of these is the Dissolution of the Elements. Could you explain the purpose of this practice, why it is undertaken, and then describe its successive stages? We can later discuss each stage in greater detail.

**00:11:01–00:12:20**

**Volunteer:** The meditation on the Dissolution of the Elements belongs to the completion stage (Dzogrim) of Highest Yoga Tantra. This is the proper context in which the practice should be understood.

**00:16:07–00:16:17**

**Interviewer:** Could you describe the successive stages in some detail and, if appropriate, tell us about your own experience at each stage, provided this does not involve teachings that should remain confidential?

**00:16:33–00:16:45**

**Volunteer:** During the stages of the Dissolution of the Elements, distinct appearances naturally arise in the practitioner's mind.

**00:22:00–00:22:10**

**Volunteer:** When earth dissolves into water, the mind experiences an appearance resembling a mirage. This is the sign associated with the dissolution of earth into water.

**00:26:30–00:26:39**

**Volunteer:** When water dissolves into fire, the appearance resembles drifting smoke.

**00:28:15–00:28:30**

**Volunteer:** When fire dissolves into wind, the appearance is like sparks flying in every direction, similar to those produced when burning dry grass during the night.

**00:29:59–00:30:22**

**Interviewer:** Words alone can hardly convey experiences of this kind to someone who has never lived through them. Nevertheless, could you describe what it feels like when these appearances arise?

**00:31:02–00:31:10**

**Volunteer:** Before describing my personal experience, we should first complete the sequence. One final stage remains: the dissolution of wind into consciousness.

**00:32:32–00:33:35**

**Volunteer:** The stages of the Dissolution of the Elements occur naturally in every sentient being at the time of death. However, without previous training, they are generally not recognized. When wind dissolves into consciousness, nothing remains as an object of appearance; only consciousness itself is present.

[The interview then turns to the subtle body, including channels (tsa) and chakras.]

**00:53:56–00:55:58**

**Dialogue:** The interviewer asks whether channels and chakras should be understood as pathways of subtle energy rather than material structures. The volunteer replies that they are difficult to describe as material entities; however, from the perspective of practice, regarding them as forms of subtle energy is not inappropriate.

**01:02:20–01:10:00**

**Volunteer:** Following the final dissolution, progressively subtler appearances arise: first a brilliant white appearance, compared to the autumn moon shining in a perfectly clear sky before dawn; then an experience known as the union of bliss and emptiness, followed by a vivid red appearance resembling clouds illuminated by the rising sun; finally, a radiant black appearance, comparable to an extraordinarily profound darkness. For practitioners who have cultivated the realization of bliss and emptiness, these stages are of great benefit. Thereafter, the Clear Light manifests.

Note: Buddhist technical terms have been translated using terminology commonly adopted in the contemporary academic literature on Vajrayāna Buddhism.

**S6. Breathing rate changes.**

No specific changes in the breathing rate were observed across PRE/ON/POST chunks.

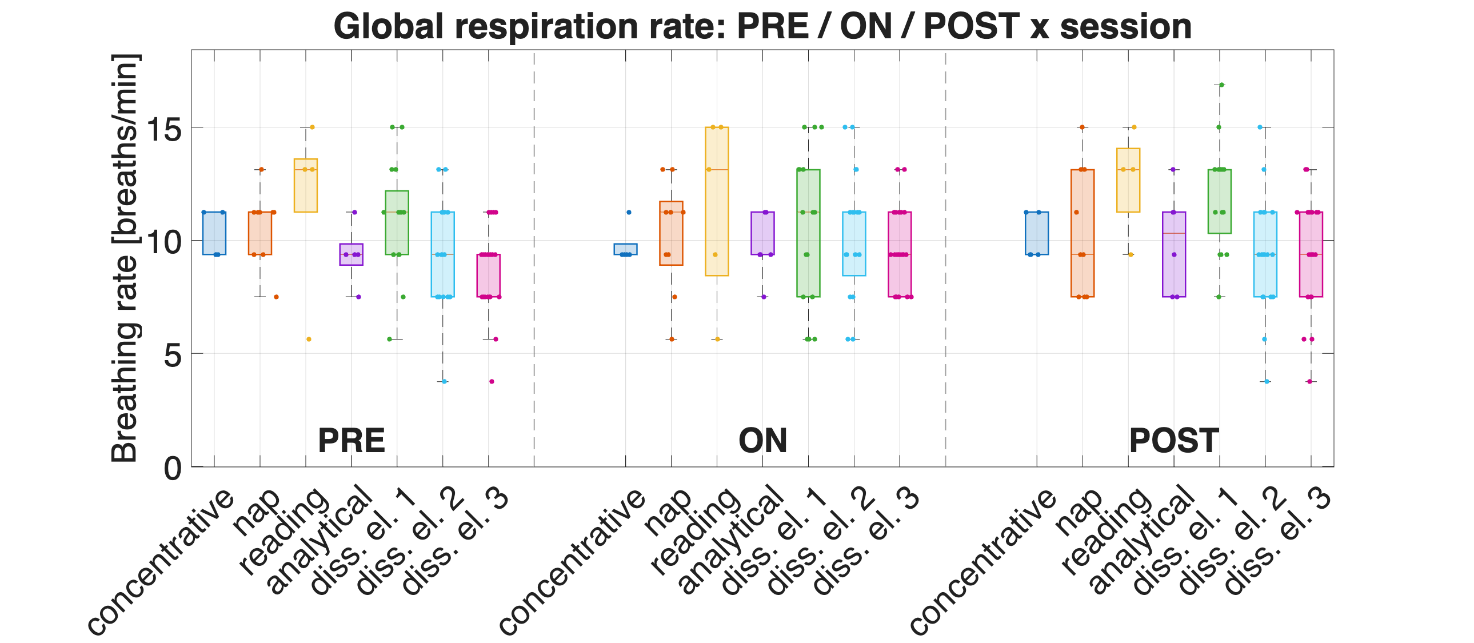

Figure S1. Boxplots of average breathing rate during matched PRE, ON and POST periods.
